# Asymmetric introgression and thermal advantage jointly drive climate-mediated lineage turnover in a mixed-ploidy reed

**DOI:** 10.64898/2026.06.02.729718

**Authors:** Lele Liu, Wenyi Sheng, Yuhui Wang, Lele Lin, Cui Wang, Huijia Song, Yaolin Guo, Weihua Guo

## Abstract

Species distribution forecasts commonly overlook intraspecific genetic variation, missing a potentially important mechanism of ecosystem change: climate-driven range shifts among lineages within a species’ native range. Here we integrate population genomic analysis of 495 individuals, multi-site common garden experiments, and species distribution modeling based on 837 occurrence records for three major genetic lineages of the foundation grass *Phragmites australis* in China. The octoploid FEAU lineage (haplotype P) exhibits superior heat tolerance (critical temperature *T_crit_* and *T_50_*) and produces significantly greater total biomass in three of four common gardens compared to the cold-adapted CN lineage (tetraploid, haplotypes O/M), which occupies a climatic niche with lower annual mean temperature (Bio1) and mean temperature of the wettest quarter (Bio8). Genomic analyses further reveal bidirectional but asymmetric introgression, with admixed individuals showing a systematic bias toward FEAU ancestry. Under the high-emission scenario (SSP5-8.5) by 2070, projected highly suitable habitat for the FEAU lineage expands by 18.6%, while the CN lineage shows a smaller relative increase. By contrast, the subtropical SW lineage (haplotypes U/I) exhibits limited and stable suitable habitat. These results demonstrate that climate change interacts with intraspecific variation among polyploidy-associated lineages, manifested through differences in thermal tolerance, biomass production, and asymmetric gene flow, to drive potential lineage replacement within a native range, a process already suggested by field observations of FEAU expansion in a plateau lake. Our findings argue for integrating evolutionary history and genetic identity into ecological forecasting to better anticipate ecosystem responses under ongoing climate warming.

## 1 Introduction

Anthropogenic climate warming is driving rapid shifts in species distributions and ecosystem composition globally, fundamentally reorganizing biodiversity (Brodie et al., 2025; Lawlor et al., 2024; Zhang et al., 2024). While species distribution models (SDMs) remain indispensable for forecasting these ecological responses, they overwhelmingly treat species as monolithic, genetically homogeneous entities (Chardon et al., 2020; Qiu et al., 2024). This assumption critically overlooks intraspecific genetic diversity, which is increasingly recognized as a primary determinant of a species adaptive capacity and vulnerability to environmental change (Cheng et al., 2021; Exposito-Alonso et al., 2022; López-Jurado et al., 2019). This knowledge gap is particularly concerning for foundation species, which disproportionately modulate community structure, habitat provision, and biogeochemical cycles (DuBois et al., 2022; Qiao et al., 2021). Consequently, ignoring their intraspecific dynamics risks missing a cryptic but profound mechanism of ecosystem reorganization: climate-driven range shifts and lineage replacement within a species native range.

The common reed, *Phragmites australis*, provides a powerful model to investigate how intraspecific variation shapes responses to global change (Eller et al., 2017; Meyerson et al., 2016). *Phragmites australis* is generally considered an allotetraploid, and hexaploids are of allopolyploid origin, whereas octoploids in Asia are most likely autopolyploid (Liu et al., 2022; Wang et al., 2024). As a nearly ubiquitous wetland foundation species, it harbors remarkable genetic and phenotypic diversity, deeply shaped by its complex evolutionary history and multiple polyploidization events (Meyerson et al., 2025; Saltonstall, 2002; Tanaka et al., 2017; Wang et al., 2024). The introduced populations of *P. australis* from Europe have shown an aggressive expansion capacity in North America (Guo et al., 2013; Saltonstall, 2002), due to smaller genome size (Pyšek et al., 2018) and greater B chromosome expansion (Wang et al., 2024; Wang et al., 2026). Similar lineage-specific dynamics may be unfolding within its native range in China, where multiple ploidy levels coexist across distinct geographic regions.

In China, which is a recognized hotspot for *P. australis* genetic diversity, three major genetic lineages occupying distinct geographic regions have been identified: the cold-adapted CN lineage (possessing O or M haplotypes) across northern China, the FEAU lineage (possessing the P haplotype, predominantly octoploid) in eastern and southern China, and the SW lineage (possessing U or I haplotypes) restricted to the subtropical southwest (Liu et al., 2022). This correspondence between genetic identity and ploidy level, initially suggested by the geographical overlap of chloroplast haplotypes (An et al., 2012) and chromosome counting-based ploidy level (C. Chen et al., 1993), has since been well established by flow cytometry (Lambertini et al., 2020), allelic number comparisons from microsatellites (Liu et al., 2022), and the alternative allele frequencies of reads mapped to the reference genome (Wang et al., 2024).

The octoploid FEAU lineage is distinguished from its tetraploid relatives not only by ploidy level but also by its distinct evolutionary history, genomic background, and geographic origin. Polyploidy has been shown in other systems to generate genetic novelty, alter gene expression, and enhance physiological stress tolerance (Bureš et al., 2024; Cheng et al., 2021; Kolář et al., 2017; Van de Peer et al., 2017), potentially pre-equipping polyploid lineages to occupy new geographical ranges and endure environmental shifts (Cheng et al., 2021; López-Jurado et al., 2019). The FEAU lineage’s superior thermal tolerance and biomass are consistent with such polyploidy-associated effects, although ploidy is correlated with, rather than experimentally separable from, the broader genetic identity of each lineage. Specifically, transcriptomic and ecophysiological evidence suggests that the octoploid FEAU lineage exhibits greater shoot height, larger leaf size, and thicker stems (K. Chen et al., 1993; Guo et al., 2025; Liu et al., 2021b, 2026; Yin et al., 2024), stronger salt tolerance and higher thermal tolerance in transcriptome (Wang et al., 2021) than its tetraploid relatives. Over the past decade, FEAU lineage of *P. australis* has expanded to Caohai Lake, a plateau lake historically not occupied by this species in China (Li et al., 2026; Ran et al., 2025). Previous work suggested asymmetric introgression between the CN and FEAU lineages, but the limited number of microsatellite markers, uneven geographic sampling, and lack of dosage correction for polyploids left its direction, magnitude, and role in climate-driven expansion insufficiently resolved (Liu et al., 2022).

Gene flow between lineages of differing ploidy can play a dual role in climate adaptation. Introgression may introduce adaptive alleles (e.g., heat tolerance loci) into a recipient lineage, facilitating its persistence under warming (Suarez-Gonzalez et al., 2018). Conversely, if introgression is asymmetric, such that one lineage’s genome is disproportionately represented in admixed populations, it can drive a gradual but systematic shift in genetic composition within the contact zone—effectively functioning as a mechanism of lineage replacement without requiring complete competitive exclusion (Bartolić et al., 2024; Zohren et al., 2016). In mixed-ploidy systems, genome dosage differences create a natural directionality in backcrossing: hybrids tend to backcross more frequently with the high-ploidy parent (Bartolić et al., 2024). In the present study, we test whether such a bias exists between the octoploid FEAU and tetraploid CN lineages and examine its consequences for future distribution under climate warming.

However, a critical question remains: how will the intraspecific differences translate into spatial biogeographic dynamics under rapid climate change? Addressing this question is essential for revealing a subtle yet potent ecological process, climate-driven lineage turnover. This process of intraspecific niche replacement represents a dynamic analogous to biological invasions but operates entirely within the native range (Paudel et al., 2025; Zhao et al., 2024). Studies that combine species distribution models with physiological or common garden experiments remain surprisingly uncommon (but see López-Jurado et al., 2019). Such integration is essential for transforming correlative SDM projections into mechanistically grounded predictions. In the present study, we adopt this integrative approach: common garden experiments directly test growth performance under controlled conditions, heat-tolerance measurements identify the specific physiological thresholds (*T*_crit_, *T*_50_) underlying lineage-specific climate responses, and SDMs project how these experimentally documented differences translate into spatial dynamics under future warming. By linking experimental data with spatial forecasting, we move beyond correlative climate matching toward a trait-based understanding of how intraspecific variation shapes species’ future distributions.

Here, we integrate population genomics, multi-site common garden experiments, and species distribution modeling to systematically investigate how historical evolutionary legacies shape climate responses among the major genetic lineages of *P. australis* in China. We explicitly test the hypothesis that the octoploid FEAU lineage possesses superior thermal tolerance and growth performance, enabling significant range expansion under future warming, whereas the other lineages will exhibit limited capacity to track climatic shifts. By linking genomic identity, physiological tolerance, gene flow, and spatial forecasting, this study provides a mechanistic test of how polyploidy-mediated intraspecific variation governs future biogeographic patterns and the structural integrity of foundation species under global change.

## 2 Materials and Methods

### 2.1 Microsatellite genotyping and population genetics

We collected 495 leaf samples of *P. australis* from 22 provinces across China **(Table S1)**. Approximately 30 mg of dried leaf tissue from each sample was placed in a 2.0-ml tube with a 5-mm glass bead, flash-frozen in liquid nitrogen, and ground using a Tissuelyser II (Qiagen) at 30 Hz. Genomic DNA was isolated using the DNAsecure Plant Kit (Tiangen). DNA integrity was checked on agarose gels, and quality and quantity were measured with a NanoDrop 2000 spectrophotometer (Thermo Scientific). Based on the established nomenclature (Saltonstall, 2002), *Phragmites australis* haplotypes were identified using sequences from the cpDNA trnT-trnL and rbcL-psaI regions; we determined the haplotypes for 202 out of 495 samples in this and previous studies.

*Phragmites australis* has a base allotetraploid genome. All 42 microsatellite markers used in this study were aligned to the *P. australis* reference genome, and each marker mapped to a single unique chromosome **(Table S2)**, confirming that each marker amplifies from one subgenome only. Therefore, in tetraploids each marker detects at most two alleles (the two homologous copies of that chromosome from the target subgenome) while the homologous region from the other subgenome is not amplified (Saltonstall, 2003). In Asia, the prevalent octoploids are most likely autopolyploid derivatives of tetraploids, carrying four homologous copies of the same chromosome and thus capable of up to four distinguishable alleles per locus (Liu et al., 2022; Wang et al., 2024). Hexaploid individuals are rare and occur primarily in contact zones, likely originating from inter-lineage hybridization (Wang et al., 2024).

In practice, the 42 selected markers very rarely produced more than four alleles in any single individual **(Table S2)**, consistent with a ploidy ceiling of octoploid. Because the exact ploidy of many samples could not be confidently assigned *a priori* (ploidy was inferred from a combination of chloroplast haplotype, geographic origin, and flow cytometry from prior studies; Lambertini et al., 2020; Liu et al., 2022), we consistently set Ploidies(mydata) <– 4 in the *polysat* R package (Clark & Jasieniuk, 2011), treating every individual as having four homologous copies. This uniform treatment is conservative: for an actual tetraploid (two copies per locus), the two unobserved “copies” are simply scored as null (missing data) in the dosage matrix; for an actual octoploid, all four detected alleles are accommodated. The allele dosage itself was estimated from high-coverage sequencing read counts (mean >5,000× per locus per sample) using the SSRSeq V1.1 pipeline (Cui et al., 2022), not inferred from allele counts alone.

Microsatellite markers were designed with SSRMMD (Gou et al., 2020) based on the transcriptome sequencing of EU and FEAU lineages of *P. australis* (Wang et al., 2021). The minimum number of repeats was 6 and 5 for trinucleotide and tetranucleotide repeats, respectively. The 64 newly developed and 5 previous (PaGT9, PaGT11, PAGT12, PAGT12, PaGT14) (Saltonstall, 2003) microsatellite markers were tested with single PCR amplification in 16 representative samples. Five newly developed markers failed to get clear bands, and the remained 64 markers were tested in the multiplex PCR. In the multiplex PCR, six marked failed to amplify, and thirteen might have variation in primer regions among samples. Therefore, a total of 58 markers were used in the microsatellite genotyping for all samples **(Table S2)**. All 58 markers were aligned to the reference genome of *P. australis* (Wang et al., 2024) using minimap2 (Li, 2021), which revealed that five markers mapped to more than one chromosome. During genotyping, eleven markers (including four of the five multi-mapping markers) were removed because more than ten samples exhibited more than four alleles per sample at these loci. Because the maximum number of distinguishable alleles expected under our ploidy model is two (tetraploid) to four (octoploid), the observation of five or more alleles in multiple individuals indicates that these markers amplify more than one genomic locus, rendering them unsuitable for dosage-based genotyping. This filtration is a quality-control step, not a ploidy assignment criterion. An additional five markers were discarded due to amplification failure in more than ten samples. One multi-mapping marker was retained owing to its ideal performance in downstream applications. Consequently, a total of 42 markers **(Table S2)** were selected for population genetic analysis. These markers are distributed across 18 out of the 25 chromosomes, including one B chromosome.

PCR amplicons (<300 bp) from each sample were pooled and tagged with 8 bp sample-specific barcodes via index primers. The barcoded products were combined into one pool for subsequent library construction. Libraries were prepared following the Illumina standard protocol and subjected to paired-end sequencing (2×150 bp) on an Illumina HiSeq 2500 platform, achieving a mean coverage of >5000× per SSR locus per sample. Raw reads were quality-checked with FastQC. An in-house Perl script named “SSRSeq count” (str_count.pl, available at https://github.com/ccoo22/SSRseq_count) was employed to process high-quality reads and generate an SSR read count table (Cui et al., 2022). Briefly, paired-end reads were merged using FLASH, and the resulting reads were aligned to *P. australis* sequences where the SSR markers were located via Blastn. The script finally produced the microsatellite read count table summarizing read counts corresponding to alleles with varying repeat numbers for each locus and sample.

Microsatellite genotyping was performed using the SSRSeq V1.1 pipeline (Cui et al., 2022; https://github.com/ccoo22/SSRseq_count). Briefly, the pipeline takes the per-locus per-sample read count table generated from high-throughput sequencing and processes it through three core steps fully described in Cui et al. (2022): (i) stutter correction, which reallocates a fraction of reads from each allele to its adjacent repeat class based on empirically estimated slip ratios; (ii) amplification bias correction, which normalizes read counts across alleles of different repeat lengths using locus-specific bias coefficients; and (iii) allele dosage calling, which selects the maximum number of alleles consistent with the specified ploidy (four in this study) and assigns integer dosages (0–4) by comparing corrected read ratios to a ploidy-adjusted threshold optimized to minimize both allelic dropout and false positives. The final output is a genotype matrix with integer allele dosages for all samples and loci, which was used directly as input to the *polysat* R package for subsequent population genetic analyses.

To delineate major evolutionary lineages, we performed principal coordinate analysis (PCoA) based on Bruvo’s genetic distance, which is robust for polyploid data and captures the primary axes of genetic differentiation without imposing a population model. To further investigate admixture and introgression among lineages, we applied Bayesian clustering in STRUCTURE, which assumes Hardy–Weinberg equilibrium within clusters and estimates individual ancestry coefficients.

We performed Principal Coordinate Analysis (PCoA) based on genetic distance using the *polysat* package (Clark & Jasieniuk, 2011) in R, which is specifically designed for analyzing polyploid microsatellite data. Since allele dosages had already been estimated based on read counts in prior steps, we did not utilize the genotype estimation functions within *polysat*. Instead, genetic distances between individuals were directly calculated using meandistance.matrix() under the default Bruvo’s distance metric. The resulting distance matrix was then subjected to classical multidimensional scaling via the cmdscale() function.

Meanwhile, we employed a Bayesian clustering approach implemented in STRUCTURE v2.3.4 (Pritchard et al., 2000) to infer individual genetic ancestry. The number of clusters (K) was tested from 1 to 10, with five independent runs per K. Each run consisted of a burn-in period of 1,000,000 steps followed by 1,000,000 MCMC iterations. The optimal K value was determined using Structure Harvester (Earl & vonHoldt, 2012). For the selected K, we aligned the replicate runs with CLUMPP (Jakobsson & Rosenberg, 2007) to obtain a consensus population structure.

Following individual ancestry inference by STRUCTURE, we classified samples into three admixture groups (pure CN, pure FEAU, and mixed) using thresholds of 80% ancestry from a single component. To visualize spatial patterns, individual data were aggregated at the provincial level and displayed on a map of China using proportional pie charts. Regional distributions of CN ancestry were examined through histograms across three geographic regions. We quantified admixture levels using an index derived from ancestry coefficients (1 – 2 × |CN – 0.5|), which was log-transformed for analysis. Linear mixed-effects models with Province as a random effect were used to assess relationships between admixture level and geographic coordinates (latitude and longitude).

A binomial test and a one-sample t-test were used to evaluate whether the proportion and mean of FEAU ancestry in admixed individuals were greater than 0.5, respectively. To infer the direction of introgression between the CN and FEAU lineages, we identified private alleles using reference groups of geographically distant pure individuals. Pure CN here reference individuals were selected from northwestern China (longitude < 100°) with STRUCTURE ancestry coefficient *Q*_CN_ > 0.9 (*n* = 70). Pure FEAU reference individuals were selected from southeastern China (longitude > 115°) with *Q*_FEAU_ > 0.9 (*n* = 44). For each of the 42 microsatellite loci, allele frequencies were calculated separately for the two reference groups. A private allele was defined as an allele with frequency > 0.05 in one reference group and < 0.01 in the opposite group. For every individual, we scored the presence or absence of private alleles from each lineage. The proportion of pure individuals (*Q* > 0.8 for the respective lineage) carrying private alleles from the opposite lineage was calculated, and binomial tests were used to assess whether these proportions were significantly greater than 0.

### 2.2 Bioclimatic niche comparisons

We compiled the geographic distribution information of major published *P. australis* haplotypes in China (O, M, P, U, I). Based on our previous microsatellite and genomic studies as well as conventional nomenclature, we designated haplotypes O and M as the CN lineage (China), P as the FEAU lineage (East Asia and Australia), and U and I as the SW lineage (southwest China, or subtropical lineage). For samples whose haplotypes were not experimentally determined in this study, lineage assignments were inferred based on results from microsatellite-based PCoA. In total, we collected 837 records, comprising 365 CN, 431 FEAU, and 41 SW lineage individuals.

All bioclimatic factors for contemporary climate conditions (1979–2013) were downloaded from WorldClim database (Fick & Hijmans, 2017) for the niche analysis. To address multicollinearity among the bioclimatic variables available, pairwise Pearson correlations were calculated **(Figure S1)**. One variable from each pair exhibiting a correlation > 0.80 was randomly excluded. The initial selection of bioclimatic factors was based on the environmental requirements of *P. australis* and previous species distribution model study of this species (Guo et al., 2013). The filtering yielded a final subset of eight key predictors for the niche comparison and further MaxEnt modeling: annual mean temperature (bio1), mean diurnal range (bio2), isothermality (bio3), temperature seasonality (bio4), mean temperature of wettest quarter (bio8), precipitation of driest month (bio14), precipitation seasonality (bio15), and precipitation of warmest quarter (bio18). Statistical differences in bioclimatic variables among lineages were assessed using Kruskal–Wallis tests with Benjamini–Hochberg (BH) *p*-value adjustment.

The environmental space of the CN and FEAU lineages was characterized using the “Characterize Environmental Space” component in the Wallace 2 platform (Kass et al., 2023). This module was employed to visualize and compare the Hutchinsonian niches of the two lineages. Specifically, a Principal Component Analysis (PCA) was performed using the Environmental Ordination module to reduce the dimensionality of the environmental data, followed by a niche overlap analysis conducted via the Niche Overlap module.

### 2.3 Common garden experiments

To investigate the growth differences and adaptive strategies between the CN and FEAU lineages, we collected growth data, including total biomass, shoot height, density, and specific leaf area (SLA), from two parallel common garden experiments: (1) A previously published common garden experiment (Song et al., 2021) conducted in 2017 across Jinnan (36.43°N, 117.45°E) and Panjin (41.20°N, 122.02°E), using CN (*n* = 11) and FEAU (*n* = 9) lineages, for which we determined the haplotype information of all samples; (2) A new common garden experiment established in 2021 across Qingdao (36.36°N, 120.69°E) and Shanghai (30.20°N, 121.29°E), with CN (*n* = 9) and FEAU (*n* = 8) lineages **(Table S3)**. Each rhizome segment (2–3 buds per segment, one segment per population) was transplanted into an individual 20 L pot, yielding one biological replicate per population per site. Although the number of populations per lineage is modest, the key inference rests on the direction and consistency of lineage differences across four climatically distinct sites rather than on the statistical significance at any single site.

In both experiments, rhizomes were propagated hydroponically and segmented to produce 5–6 segments per population (each with 2–3 buds), which were then transplanted individually into 20 L pots containing a standardized soil substrate. Identical populations were planted simultaneously at both sites under rigorously controlled conditions, with uniform substrate, container size, fertilization, and pesticide application, thereby isolating ambient climate as the sole environmental variable. Garden management was synchronized across sites throughout the growing season, including bi-weekly irrigation (adjusted for ambient precipitation), monthly application of balanced fertilizer from June to September, and standardized pest and disease control.

Measurement protocols were consistent across both gardens. Plant height was defined as the distance from the soil surface to the tallest inflorescence or, in its absence, the highest leaf tip. For each pot, the heights of the three tallest shoots were measured and averaged. Five fully expanded leaves (from nodes 3–5) per pot were randomly selected, placed in labeled plastic bags, and rehydrated in darkness for 12 hours prior to measuring saturated fresh mass. Leaf areas were determined from scanned images using ImageJ software calibrated with a millimeter scale. All leaf samples were then oven-dried at 80 °C for 48 hours and weighed. Aboveground biomass was harvested by cutting shoots at soil level, placed in paper bags, and dried at 80 °C until constant mass was achieved. Total belowground dry mass was estimated by extrapolating the fresh-to-dry mass ratio obtained from a subsample to the entire root system.

For each trait separately, we calculated the mean and standard error (SE) for each lineage within each garden. To assess the statistical significance of differences between the two lineages within each garden, pairwise Student’s *t*-tests were performed independently for each garden-trait combination. No correction for multiple testing was applied, as each common garden was considered an independent experimental environment.

### 2.4 Heat tolerance measurement

Heat tolerance was assessed on a sunny morning after approximately three months of cultivation. For each of the 12 *P. australis* genotypes (6 FEAU and 6 CN) grown in a common garden (36.38°N, 118.90°E) located in Weifang, Shandong Province, China in 2024, the third or fourth fully expanded leaves were selected. After excision, the leaves were wrapped in moist tissue paper, placed in ziplock bags with ice packs, and transported to the laboratory under dark conditions to minimize transpiration. In the laboratory, the leaves were cut into segments approximately 1 cm in length. These segments were then wrapped in wet filter paper, sealed in plastic bags, and subjected to heat treatment in a water bath. The temperature regimen consisted of sequential exposure to 25 °C (ambient control), 38 °C, 40 °C, 42 °C, 44 °C, 46 °C, 48 °C, 50°C, 52 °C, 54 °C, 57 °C, and 60 °C. Following heat treatment, all samples were kept in moist, dark conditions for 24 hours. After dark adaptation, the maximum quantum yield of photosystem II (Fv/Fm) was measured using a pulse-amplitude modulation (PAM) fluorometer (PAM-2500, Walz, Effeltrich, Germany). Each temperature treatment included five biological replicates.

Parameters of heat tolerance (*T_crit_*, *T_50_*, and *T_95_*) were derived from a logistic decay model that describes the relationship between Fv/Fm and temperature. Parameter estimates were obtained through nonlinear least-squares regression using the *nls* routine in R. Mean values for each parameter were calculated using bootstrap resampling, implemented within a custom R function (psiiht), which also provided optional diagnostic plotting (Perez et al., 2021). The critical temperature (*T_crit_*) was identified as the temperature at which the slope of the Fv/Fm decline reached 15% of the maximum negative slope observed during heating, with the additional constraint that this point must occur below the temperature causing 50% inhibition. The temperature of 50% inhibition (*T_50_*) was defined as the point at which Fv/Fm decreased to half of the value measured at the control temperature. Similarly, the temperature of 95% inhibition (*T_95_*) corresponds to a 95% reduction in Fv/Fm relative to the control level. All three parameters were compared between CN and FEAU lineages using a *t*-test.

### 2.5 Species distribution models

To assess the impact of climatic changes on the distribution of *P. australis* lineages, species distribution models were developed using eight selected bioclimatic variables. Potential geographic distributions were projected using the maximum entropy algorithm (MaxEnt) within Wallace (Kass et al., 2023). MaxEnt models species distributions by comparing environmental conditions at occurrence sites with those across the background environment. The model was calibrated with 10,000 randomly generated background points from the study region. Regularization multipliers were tested ranging from 0.5 to 5.0 at intervals of 0.5, and all feature class combinations (Linear, Quadratic, Hinge, Product) were examined. Model selection followed a random k-fold cross-validation approach (*k* = 4), with the best model chosen based on the lowest corrected Akaike Information Criterion (ΔAICc).

For projecting future distributions across both native and introduced ranges, bioclimatic variables at 5-arcmin resolution were obtained from WorldClim. Future projections for the period 2061–2080 incorporated two representative climate scenarios: the low-emission pathway SSP1-2.6 and the high-emission pathway SSP5-8.5 (IPCC, 2021). Future climate projections were derived from the Centre National de Recherches Météorologiques Earth System Model 2-1 (CNRM-ESM2-1) (Séférian et al., 2019).

## 3 Results

### 3.1 Genetic structure and lineage delimitation

Given that STRUCTURE showed optimal *K* = 2 **(Figure S2)** and failed to fully resolve the SW lineage **(Figure 1A)**, we used PCoA for lineage delineation **(Figure 1B)** and STRUCTURE for admixture quantification **(Figure 2)**. This complementary approach allows us to distinguish between evolutionary lineage identity and contemporary gene flow.

**Figure 1.**
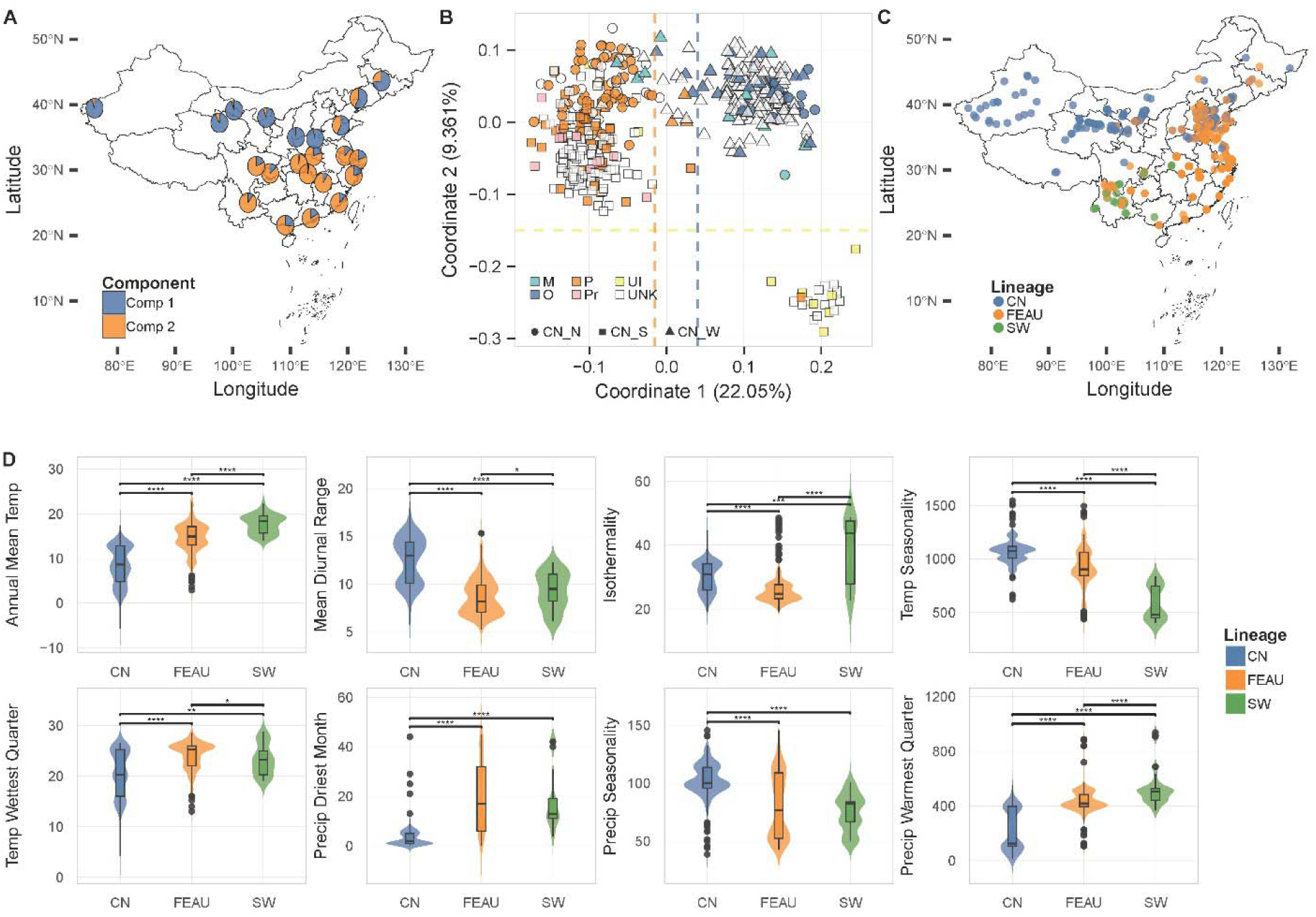
Evolutionary lineages of *Phragmites australis* in China. **(A)** Bayesian clustering analysis based on 42 nuclear microsatellites at the optimal genetic cluster number *K* = 2. **(B)** Principal coordinate analysis (PCoA) based on Bruvo’s genetic distance. Colors indicate chloroplast haplotypes, where P_r denotes P-related haplotypes and “Unknown” refers to samples without haplotype data. Shapes denote geographical origins: circles (northern China, CN_N), squares (southern China, CN_S), and triangles (northwestern China, CN_W). **(C)** Geographical distribution of sampling locations for each genetic lineage by PCoA or/and chloroplast haplotypes. **(D)** Bioclimatic variable differences among lineages. Bio1, annual mean temperature; bio2, mean diurnal range; bio3, isothermality; bio4, temperature seasonality; bio8, mean temperature of wettest quarter; bio14, precipitation of driest month; bio15, precipitation seasonality; bio18, precipitation of warmest quarter. Significance levels are indicated as: * *p* < 0.05, ** *p* < 0.01, *** *p* < 0.001.

**Figure 2.**
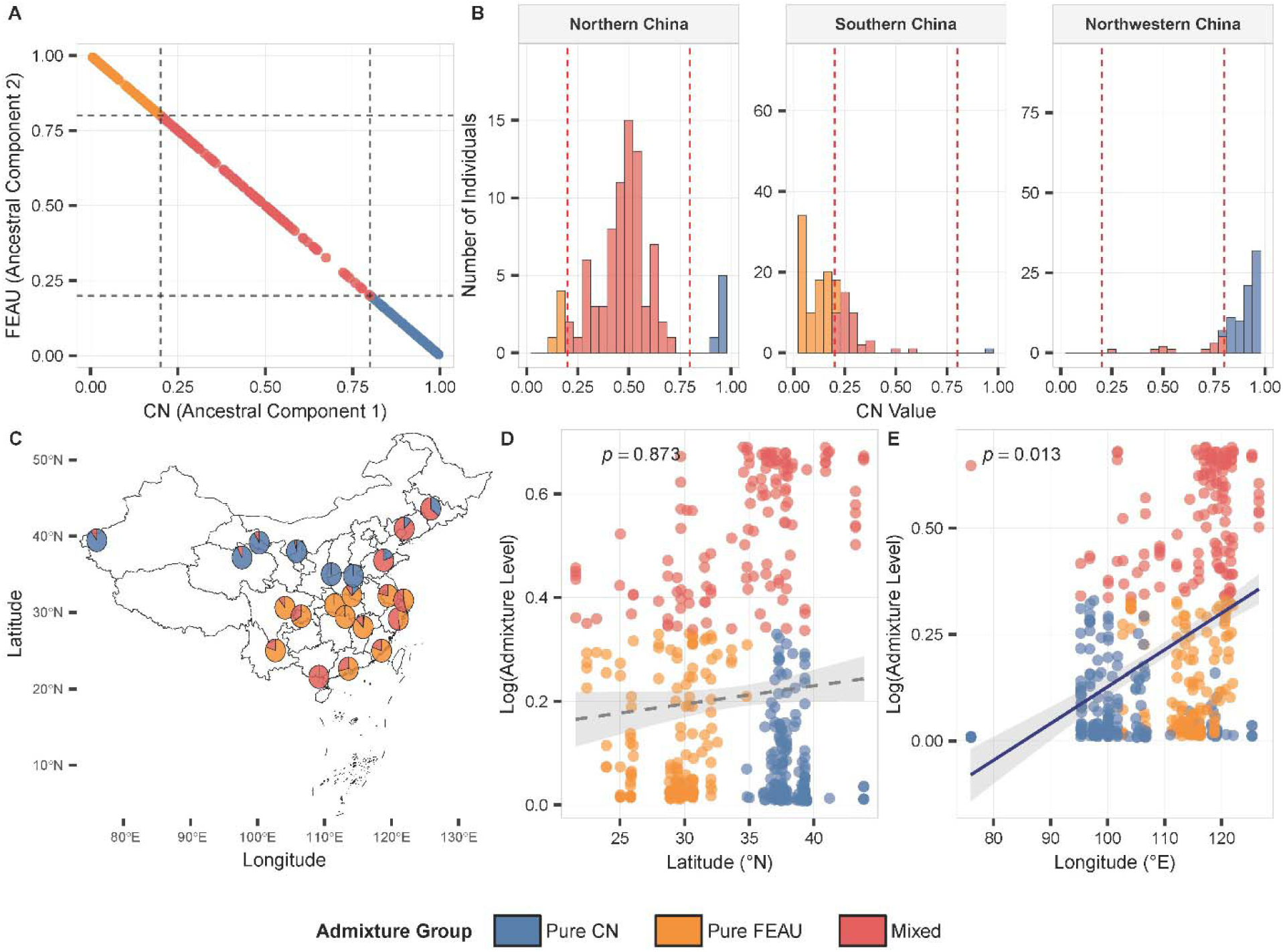
Admixture and introgression between CN and FEAU lineages revealed by STRUCTURE. **(A)** Scatter plot of individual ancestry coefficients for the two ancestral components (CN and FEAU). Dashed lines indicate the 0.2 and 0.8 thresholds used to define pure and admixed individuals. Points are colored by admixture group. STRUCTURE is used here to detect hybridization, not to define lineages (see Figure 1). **(B)** Histograms of CN ancestry values across three geographic regions: Northern China, Southern China, and Northwestern China. **(C)** Geographic distribution of admixture groups across Chinese provinces. Pie charts show the relative proportions of pure CN (blue), pure FEAU (orange), and mixed (red) individuals in each province. **(D)** Relationship between latitude and log-transformed admixture level. Points are colored by admixture group. The regression line (solid if *p* < 0.05, dashed if *p* > 0.05) and *p*-value (from linear mixed-effects model with Province as random effect) are shown. The admixture level for each individual was calculated as 1 – 2 × |CN−0.5|, where CN is the ancestry coefficient of the CN lineage (range 0 – 1). This value approaches 1 when CN = 0.5 (high admixture) and approaches 0 when CN is near 0 or 1 (near-pure ancestry). To improve visualization, we plotted log (admixture level + 1), which ranges from 0 (pure) to log (2) ≈ 0.69 (maximally admixed). **(E)** Relationship between longitude and log-transformed admixture level. Visualization follows the same conventions as panel D. Pure individuals are defined as having > 80% ancestry from one component; individuals with 20-80% ancestry from each component are classified as mixed.

The PCoA clearly resolved three major genetic clusters, along with a few introgressive and nuclear-chloroplast discordant samples **(Figure 1B)**. Coordinate 1, explaining 22.05% of the variance, separated the FEAU lineage (haplotype P and related haplotypes such as AS) from the CN lineage (haplotypes O and M). Coordinate 2, which explained 9.36% of the variance, distinguished the SW lineage (haplotypes U and I).

For the Bayesian clustering analysis in STRUCTURE, at *K* = 2, two genetic clusters were identified, distributed across northern and southern China, respectively **(Figure 1A)**, corresponding to the previously reported CN and FEAU lineage. When K was increased to 3, a southwestern genetic component was further differentiated **(Figure S3)**.

Admixture between CN and FEAU lineages was widespread across China, with a general predominance of FEAU ancestry **(Figure 2A)**. The spatial distribution of admixture groups, however, exhibited distinct regional patterns **(Figure 2B–E)**. Southern China was predominantly occupied by pure FEAU and mixed individuals, northwestern China was dominated by pure CN with a limited presence of mixed types, while northern China was primarily characterized by admixed individuals alongside both pure FEAU and CN lineages **(Figure 2B, C)**. These geographical gradients resulted in a significant longitudinal effect on admixture levels (*p* = 0.013), whereas no significant latitudinal pattern was observed (*p* = 0.873) **(Figure 2D, E)**.

Among the 131 individuals classified as admixed (*Q*_CN_ and *Q*_FEAU_ both between 0.2 and 0.8), 80 individuals (61.1%) had a FEAU ancestry proportion greater than 0.5 (0.569 ± 0.16). Both the proportion (binomial test, *p* = 0.007) and the mean (one-sample *t*-test, *t* = 4.89, *df* = 130, *p* < 0.001) were significantly greater than 0.5, indicating a systematic bias toward FEAU ancestry in admixed individuals. Using reference groups of 70 pure CN individuals and 44 pure FEAU individuals, we identified 16 CN-private alleles and 19 FEAU-private alleles across the 42 microsatellite loci. Among 194 pure CN individuals (*Q*_CN_ > 0.8), 56 individuals (28.9%) carried at least one FEAU-private allele (binomial test, *p* < 0.001). Among 168 pure FEAU individuals (*Q*_FEAU_ > 0.8), 77 individuals (45.8%) carried at least one CN-private allele (*p* < 0.001). These results demonstrate bidirectional gene flow between the two lineages, with CN → FEAU introgression being more prevalent at the level of private alleles.

### 3.2 Differentiation of bioclimatic niche among lineages

Significant differences were observed among lineages across all eight bioclimatic variables analyzed **(Figure 1C)**. For bio1, bio3, bio8, and bio18, the FEAU lineage exhibited values significantly higher than those of CN but lower than those of SW. In contrast, for bio2 and bio4, values in FEAU were significantly lower than in CN but higher than in SW. Meanwhile, CN showed significantly lower values than both FEAU and SW in bio14, but higher values in bio15 **(Figure 1D)**.

Following environmental PCA **(Figure S4)**, niche similarity tests revealed a statistically significant moderate overlap between the CN and FEAU lineages **(Figure S5)** (Schoener’s *D* = 0.48, *p* = 0.01). Specifically, 16% of the environmental space was occupied exclusively by the CN lineage, while 40% was unique to the FEAU lineage. Approximately 60% of the environmental space was shared by both lineages, indicating considerable common habitat suitability despite their distinct ecological characteristics.

### 3.3 Differentiation of growth performance between two lineages

The four common gardens are all located in eastern China where the CN and FEAU lineages co-occur, with mean annual temperature (Bio1) ranging from 9.3 °C (Panjin) to 16.8 °C (Shanghai) **(Figure 3A)**. The total biomass of the FEAU lineage was significantly greater than that of the CN lineage in Jinan, Panjin, and Qingdao, but no significant difference was observed in Shanghai **(Figure 3B)**. Similarly, plant height was significantly greater in the FEAU lineage than in the CN lineage in Jinan, but not in the other common gardens (**Figure 3C**). Effect sizes for total biomass were large in Jinan (Cohen’s *d* = 1.10), Panjin (*d* = 1.12), and Qingdao (*d* = 1.43), but negligible in Shanghai (*d* = 0.37), confirming that the lineage differences, where present, are biologically substantial.

**Figure 3.**
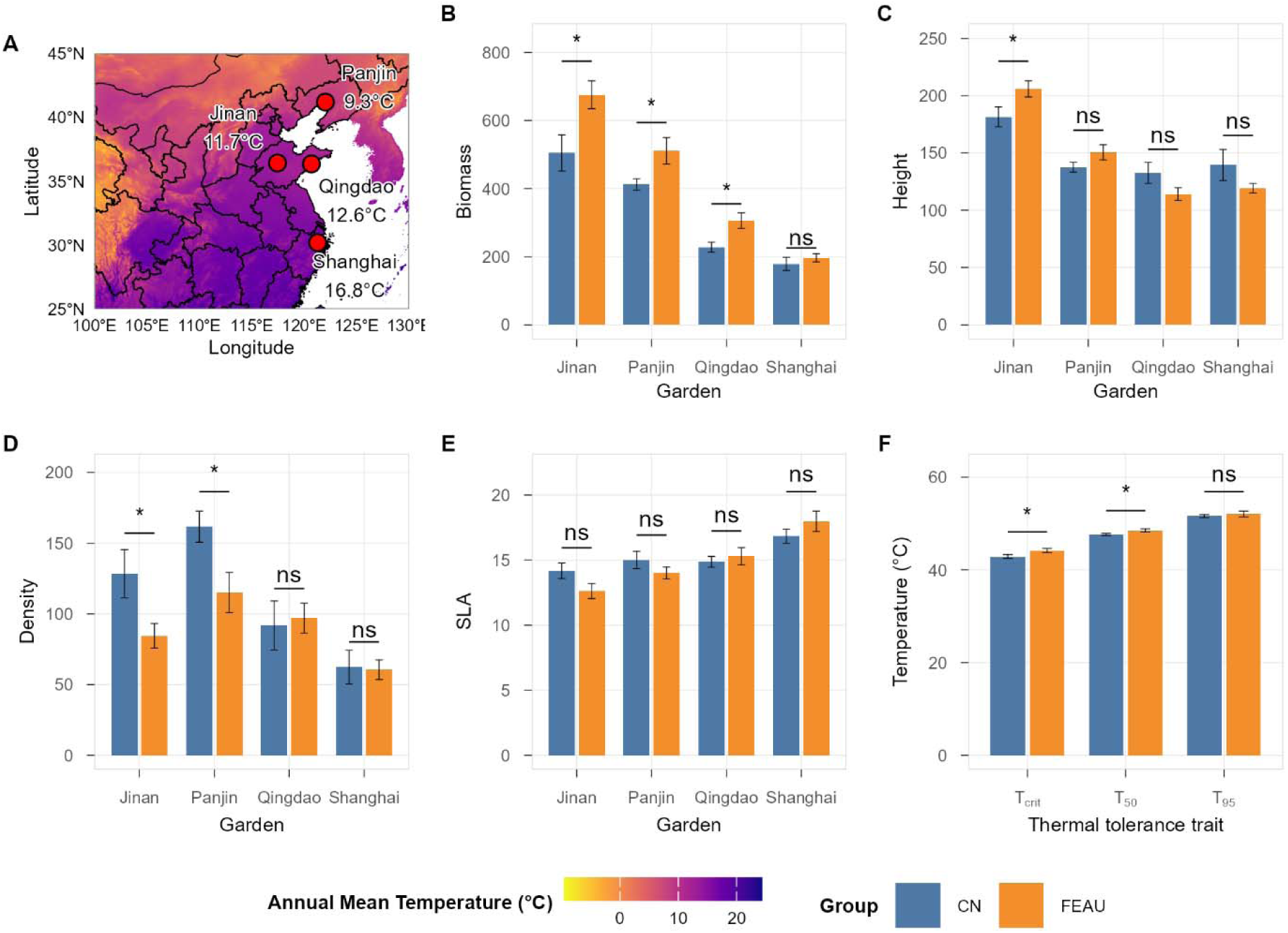
Comparative growth performance and heat tolerance of the CN and FEAU lineages of *Phragmites australis* across common garden environments. **(A)** Geographic locations of the four common garden sites (Panjin, Jinan, Qingdao, Shanghai) overlaid on the mean annual temperature (Bio1) raster map of eastern China. Color gradient represents annual mean temperature (°C). **(B-E)** Comparison of four growth traits between lineages across four common garden locations: **(B)** Total Biomass, **(C)** Shoot Height, **(D)** Density, and **(E)** Specific Leaf Area (SLA). Bar heights represent mean values, and error bars indicate the standard error of the mean. **(F)** Comparison of key heat tolerance parameters (*T_crit_*, *T_50_* and *T_95_*) between the CN and FEAU lineages. Significance levels from pairwise *t* tests within each garden **(B-E)** and unpraised *t* tests for heat tolerance parameters **(F)** are denoted by asterisks: * *p* < 0.05, ** *p* < 0.01, *** *p* < 0.001; “ns” indicates not significant.

In contrast, the CN lineage exhibited significantly higher density than the FEAU lineage in Jinan and Panjin, while no significant differences were detected in Qingdao or Shanghai (**Figure 3D**). For specific leaf area (SLA), no significant differences were found between the two lineages in any of the common gardens (**Figure 3E**).

### 3.4 Heat tolerance of two lineages

The heat tolerance parameters *T_crit_*, *T_50_*, and *T_95_* were compared between CN and FEAU lineages (**Figure 3F**; **Figure S6; Table S4)**. The mean *T_crit_* for the CN group was 42.9 ± 1.04 °C, which was significantly lower than that of the FEAU group (44.2 ± 0.96 °C; *p* < 0.05). Similarly, a significant difference was observed in *T_50_*, with the CN group exhibiting a mean of 47.7 ± 0.56 °C compared to 48.5 ± 0.78 °C in the FEAU group (*p* < 0.05). In contrast, no significant difference was detected in *T_95_* between the two groups; the mean values were 51.6 ± 0.83 °C for CN and 52.1 ± 1.36 °C for FEAU.

### 3.5. Potential distribution projection of three lineages

The optimal model configurations and performance metrics varied among the three lineages **(Table S5)**. The CN lineage achieved the best fit with the LQHP feature class and a regularization multiplier of 0.5, resulting in high predictive accuracy (AUCtrain = 0.947) and excellent model stability (CBItrain = 0.973). This model was strongly supported (AICc weight ≈ 1.000). The FEAU lineage performed best under the LQH feature class and RM = 0.5, also showing high training AUC (0.946) and good validation performance (AUCval = 0.898). In contrast, the SW lineage was best modelled using a simpler LQ feature class with RM = 1.0, yielding the highest training AUC (0.965) but lower complexity (7 coefficients). All models demonstrated high predictive performance, with training AUC values exceeding 0.94.

Model projections indicated notable shifts in habitat suitability under future climate scenarios **(Figure 4; Figure S7; Table S6)**. For the CN lineage, the high-emission scenario (SSP5-8.5) resulted in a marked expansion of high-suitability areas to 16.5%, compared to 9.4% under current conditions. In contrast, the FEAU lineage showed the most pronounced response under SSP5-8.5, with high-suitability habitat increasing substantially to 25.2%, while low-suitability areas contracted to 56.7%. The SW lineage remained relatively stable across projections, with the majority of habitats consistently classified as low suitability, though a slight increase in high and medium suitability was observed under SSP5-8.5. These results suggest divergent responses to climate change among lineages, with FEAU exhibiting the greatest potential for range expansion under high-emission scenarios.

**Figure 4.**
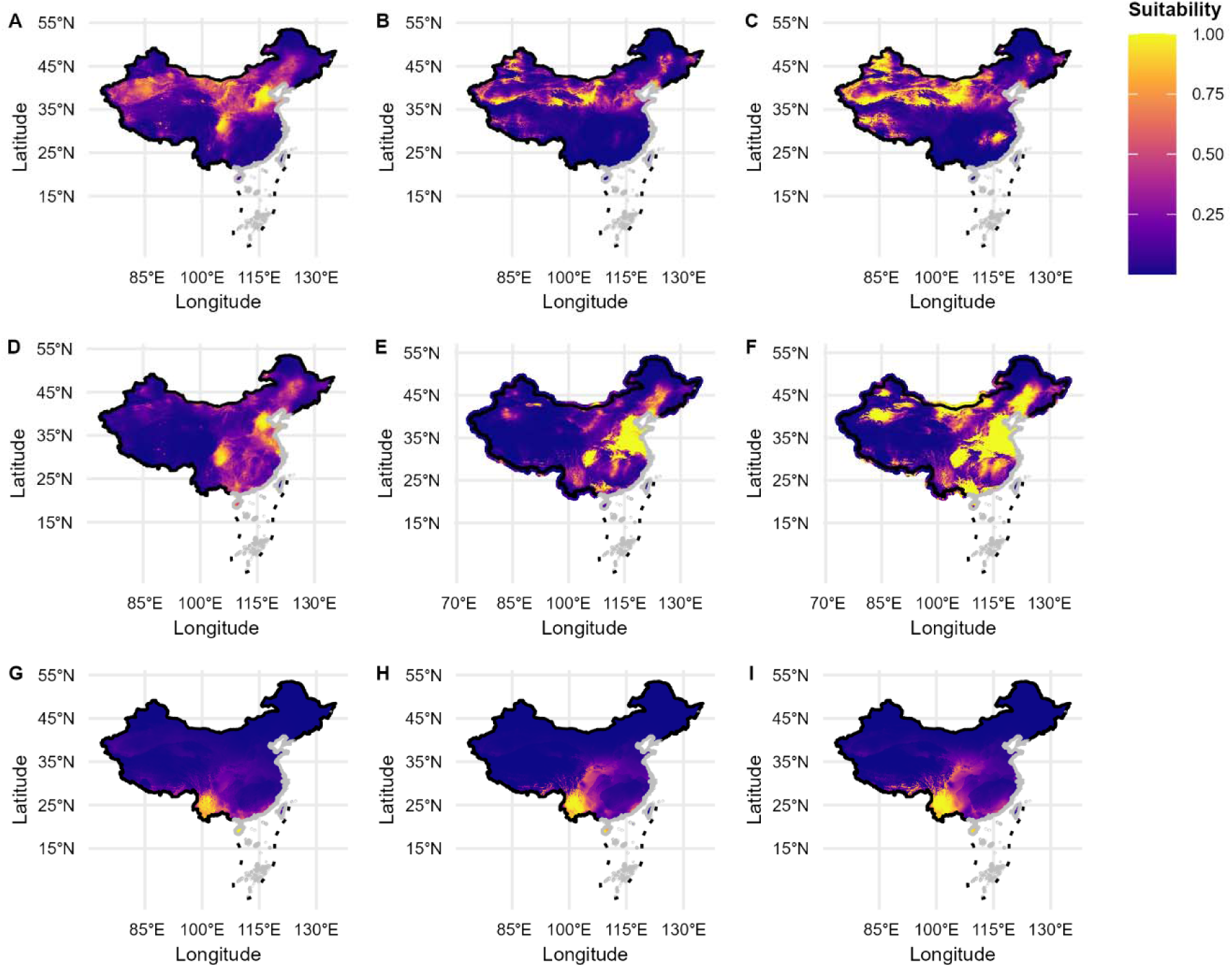
Predicted potential distribution of three *Phragmites australis* lineages (CN, FEAU, and SW) in China under current and future climate scenarios. The first row (A, B, C) shows the results for the CN lineage, the second row (D, E, F) for the FEAU lineage, and the third row (G, H, I) for the SW lineage. Columns represent different time periods: current distribution (A, D, G), and projected distributions for 2061–2080 under the low-emission scenario SSP1-2.6 (B, E, H) and the high-emission scenario SSP5-8.5 (C, F, I). Predictions were generated using the Maximum Entropy (MaxEnt) model. The depth of color (color intensity) corresponds to the level of habitat suitability, ranging from dark blue (unsuitable) to bright yellow (highly suitable).

## 4 Discussion

### 4.1 Ecophysiological basis of thermal tolerance and range expansion

Our findings demonstrate that the octoploid FEAU lineage of *P. australis* possesses greater heat tolerance and biomass production than the tetraploid CN lineage. Under a high emission scenario (SSP5 8.5), the projected suitable habitat for the FEAU lineage expands by 18.6%, while the CN lineage exhibits a much smaller relative increase. Several non-mutually-exclusive mechanisms could explain these lineage-level differences, including increased gene dosage from whole-genome duplication, divergent selection histories, and/or standing genetic variation in thermal tolerance loci unlinked to ploidy (Bures et al., 2024; Cheng et al., 2021; Van de Peer et al., 2017). Our data cannot fully partition these factors, but the strong association between lineage identity and both physiological performance and projected range dynamics highlights the importance of incorporating intraspecific lineage information into ecological forecasts, regardless of the ultimate causal mechanism.

The superior heat tolerance of the FEAU lineage, characterized by an approximately 1.3 °C higher *T_crit_* and 0.8 °C higher *T_50_*, aligns with evidence that whole genome duplication can enhance stress tolerance through gene dosage effects, subfunctionalization, or epigenetic modifications (Van de Peer et al., 2017). In *P. australis*, transcriptomic comparisons have revealed that octoploids upregulate heat shock proteins and maintain photosystem II integrity under elevated temperatures (Wang et al., 2021). Similar patterns occur in other polyploid systems, such as *Dianthus broteri* (López-Jurado et al., 2019) and *Fragaria* (Wei et al., 2020), although this advantage can be context dependent (Kolář et al., 2017).

A 1.3 °C difference in *T_crit_* could be critical when summer temperatures approach or exceed a lineage’s photosynthetic thermal thresholds. Mean annual air temperature over China has risen by more than 1 °C, with more frequent and hotter summer days (Wei et al., 2023). In eastern China, extreme heat events are becoming more frequent and intense, with peak summer temperatures increasingly approaching the critical thresholds (42.9 °C for CN, 44.2 °C for FEAU) identified here (Zhou et al., 2024). Direct evidence linking PSII heat tolerance to survival in wetland plants remains limited (Perez et al., 2021), yet the functional consequence is clear: a lineage that maintains photosynthesis at 44 °C rather than ceasing at 43 °C will sustain carbon assimilation longer during heatwaves. Unlike woody plants in drylands, where heatwave mortality often results from hydraulic failure (Breshears et al., 2021), *P. australis* in wetlands is unlikely to face severe water limitation; direct thermal damage to the photosynthetic apparatus is the primary constraint. Even marginal differences in thermal tolerance can be amplified through positive feedback: prolonged carbon assimilation leads to greater biomass, which in turn enhances competitive ability and accelerates range expansion (Cheng et al., 2021). Therefore, the consistent physiological differences documented here are likely to translate into meaningful demographic advantages under the intensifying heat regimes expected across eastern China.

### 4.2 Niche overlap, admixture, and the trajectory of lineage replacement

Bayesian clustering and principal coordinate analysis resolved three distinct genetic lineages, revealing a broad zone of admixture between the FEAU and CN lineages in northern China. A substantial proportion of individuals exhibited mixed ancestry, with admixture levels increasing significantly with longitude but not latitude, consistent with a previous study using limited microsatellite markers (Liu et al., 2022). This east–west gradient suggests ongoing gene flow, which may be facilitated by both natural and human-mediated dispersal. Eastward water dispersal via the Yellow River (Liu et al., 2021a, 2021b) and historical introductions of *P. australis* germplasm for reed production in areas such as the Panjin region (Brix et al., 2014) likely contribute to this pattern. However, the persistence of pure CN and pure FEAU individuals within admixed populations implies that reproductive barriers or selection against hybrids exist, potentially driven by ploidy differences (tetraploid versus octoploid) or local adaptation.

The moderate environmental niche overlap between CN and FEAU (Schoener’s *D* = 0.48) indicates that approximately 60% of their environmental space is shared. Under current climatic conditions, parapatric coexistence and the observed admixture patterns may be maintained by dispersal limitations or competitive trade-offs (Guo et al., 2024; Liu et al., 2026). For example, the CN lineage’s higher shoot density at northern sites may contribute to its persistence in cooler regions, potentially offsetting the demographic disadvantages of its lower heat tolerance and morphological size. However, coexistence theory predicts that when two lineages share a large fraction of environmental space and one possesses a consistent performance advantage, competitive exclusion is likely to occur in a changing environment (Pastore et al., 2021).

Furthermore, the octoploid FEAU lineage likely possesses greater genomic compatibility than the tetraploid CN lineage, leading to asymmetric introgression: hybrids are expected to backcross more frequently with the high-ploidy lineage (Bartolić et al., 2024; Han et al., 2015; Liu et al., 2022; Zohren et al., 2016), gradually shifting the genetic composition of admixed populations toward the FEAU genome. Supporting this hypothesis, private allele analysis revealed bidirectional gene flow, with 28.9% of pure CN individuals carrying FEAU-private alleles and 45.8% of pure FEAU individuals carrying CN-private alleles. Crucially, admixed individuals showed a significant bias toward FEAU ancestry (61.1%), consistent with asymmetric backcrossing favoring the octoploid genome.

Our common garden results corroborate this trajectory: the FEAU lineage produced significantly greater biomass than CN in three gardens (Jinan, Panjin, and Qingdao), but not in the warmest (Shanghai). This exception may reflect other limiting factors in Shanghai or indicate that FEAU’s thermal advantage diminishes near its upper physiological limits. Because the two experiments differed in starting material and growth duration, we focus only on the direction and significance of lineage differences within each site, not on absolute biomass across gardens.

As warming progresses, the competitive balance is expected to shift decisively toward FEAU. The current admixture zone may act as a moving hybrid front, with FEAU gradually displacing CN in warmer regions while potentially maintaining a stable hybrid zone in transitional climates. This projected lineage replacement differs fundamentally from the non-native invasion of *P. australis* in North America (Saltonstall, 2002), where an introduced European lineage spread into novel territory; in China, the expansion involves only native genotypes. Nevertheless, both processes threaten to erode unique genetic diversity and reduce the species’ adaptive capacity.

From an evolutionary standpoint, asymmetric introgression can erode the genetic distinctiveness of the minority lineage (CN) while enriching the majority lineage (FEAU) with alleles that may have been locally adapted in the CN genomic background. This could reduce the species’ overall evolutionary potential, even if the FEAU lineage itself thrives, because cold-adapted alleles from the CN lineage, which may be valuable under future climate volatility (including extreme cold events), risk being diluted or lost (Exposito-Alonso et al., 2022). The directionality of introgression is also not fixed; it could shift if environmental conditions alter hybrid fitness or if the demographic balance between lineages changes. Long-term genomic monitoring of the CN–FEAU contact zone will be essential to determine whether the asymmetric gene flow documented here represents a transient phase or a persistent trajectory toward genomic homogenization.

### 4.3 Vulnerability of the SW lineage and potential ecosystem consequences

The SW lineage, with only 41 occurrence records and a restricted distribution, occupies a distinct climatic niche and is projected to remain spatially limited under future scenarios. This stability likely reflects a narrow fundamental niche rather than resilience. Field observations of FEAU encroachment into Caohai Lake, a site historically not occupied by *P. australis* (Li et al., 2026; Ran et al., 2025), echo patterns seen in other systems where warm adapted lineages displace cold adapted ones (Cheng et al., 2021; Paudel et al., 2025). Although not yet quantified, this trend merits urgent monitoring. The SW lineage should be considered a high priority conservation target, and its habitats must be carefully managed to minimize the inadvertent introduction of FEAU propagules (Tanaka et al., 2017).

The potential replacement of the CN lineage by the FEAU lineage, if realized, could have ecosystem-level consequences beyond genetic composition. As a foundation species, *P. australis* modulates wetland structure and biogeochemical cycles. The FEAU lineage’s tendency toward greater biomass but lower shoot density may alter light availability, sediment dynamics, and habitat complexity for invertebrates (Yan et al., 2021). However, these ecosystem consequences remain speculative. Direct measurements of decomposition rates, greenhouse gas fluxes, and faunal communities across monospecific stands of each lineage are required to test these hypotheses.

### 4.4 Limitations, future directions, and conservation implications

While our SDM projections provide a robust framework for assessing vulnerability, they inherently assume unlimited dispersal and an absence of novel biotic interactions (Di Cola et al., 2017; Phillips et al., 2017). The natural expansion of the FEAU lineage will ultimately depend on propagule dispersal distances (seeds and rhizomes) and landscape barriers (Liu et al., 2021a).

The current admixture zone in northern China confirms that gene flow is active, but the actual rate of spread may lag model projections. Future work should integrate mechanistic dispersal models and long-term field monitoring of contact zones to validate these spatial predictions. Moreover, precipitation changes and land use conversion were not fully accounted for, both of which could alter habitat suitability across regions (Guo et al., 2013).

We also acknowledge that the common garden experiments, while replicated across four climatically distinct sites, involved a limited number of populations per lineage (9–11 CN and 8–9 FEAU), which constrains our ability to fully separate lineage-level effects from population-level variation. The consistent direction of biomass differences across three of four sites, supported by large effect sizes, nonetheless provides robust evidence for a lineage-level performance advantage that merits further confirmation with a larger, more geographically representative panel of populations.

Crucially, ploidy was not experimentally manipulated in this study; it is inherently confounded with the distinct evolutionary history and genomic background of each lineage. While the observed thermal tolerance and biomass differences are consistently associated with the octoploid FEAU lineage, we cannot formally exclude the possibility that these traits are driven by genetic factors independent of ploidy *per se*. Future studies using experimental approaches that can partition ploidy effects from lineage-specific genetic effects, such as common gardens with synthetic polyploids or transcriptomic analyses comparing gene expression dosage responses, are needed to strengthen causal inference (Wei et al., 2020).

Similarly, the common garden results should be interpreted as lineage-associated rather than ploidy-causal performance differences. The potential role of admixture in facilitating the adaptive introgression of heat tolerance alleles also warrants deeper investigation (Suarez-Gonzalez et al., 2018).

We also recognize that our SDM approach, while disaggregating the species into three major genetic lineages, still treats each lineage as a homogeneous entity. This simplification parallels—albeit at a finer scale—the species-as-monolith assumption that we critique in the Introduction. Within-lineage variation in thermal tolerance, growth, and dispersal capacity is plausible, particularly given the broad geographic ranges of the CN and FEAU lineages. By modelling each lineage as a uniform group, our projections may overestimate the precision of range forecasts and underestimate the evolutionary potential of standing variation within lineages (Chardon et al., 2020). Future frameworks that incorporate trait variation at multiple hierarchical levels (population, lineage, ploidy) will be necessary to capture both the adaptive potential and the ecological constraints that shape species’ responses to climate change.

Looking beyond the present study, we envision three complementary directions that build upon our current findings. First, expanding common garden experiments to include admixed individuals would test whether introgressed genomic blocks confer fitness advantages under thermal stress. Second, whole-genome resequencing coupled with selection scans and genotype-environment association analyses could pinpoint adaptive loci and reveal whether heat-tolerance alleles are preferentially transferred via asymmetric introgression. Third, integrating transcriptomic profiling with phenotypic measurements would help disentangle ploidy effects from genetic background. Together, these directions form an integrated framework that moves from the correlative patterns reported here toward mechanistic understanding.

The projected replacement of a cold adapted native lineage by a warm adapted native lineage presents a nuanced conservation challenge. Unlike classical biological invasions, no non-native species is involved; nevertheless, the loss of unique regional gene pools (CN and SW) may erode the species’ overall adaptive capacity to future environmental volatility (Exposito-Alonso et al., 2022). To safeguard evolutionary resilience, we recommend three practical measures: using strictly locally sourced germplasm in wetland restoration to prevent the human mediated spread of the FEAU lineage; protecting established hybrid zones as reservoirs of genetic diversity; and establishing long-term monitoring networks in geographic contact zones to detect lineage turnover early (Brodie et al., 2025; Lawlor et al., 2024; Vranken et al., 2021).

## Conclusions

The integration of population genomics, common garden experiments, and species distribution modeling reveals three interacting mechanisms that jointly drive lineage turnover under climate warming. First, the octoploid FEAU lineage possesses superior thermal tolerance (1.3 °C higher *T_crit_*) compared to the tetraploid CN lineage. Second, the octoploid lineage exhibits intrinsically greater biomass production, a trait that is likely an inherent consequence of polyploidy and further reinforced by its thermal tolerance, together enhancing its competitive ability. Third, asymmetric introgression, resulting from preferential backcrossing of hybrids with the octoploid FEAU lineage, progressively shifts the genomic composition of admixed populations toward FEAU. These mechanisms, coupled with moderate niche overlap and ongoing gene flow, strongly suggest a trajectory of gradual lineage replacement across eastern China. This prediction is supported by initial field observations of FEAU expansion in a plateau lake (Li et al., 2026), but decadal monitoring and dispersal constraints are needed for validation. Incorporating evolutionary history and functional genetic identity into ecological forecasting remains essential as climate change reshapes ecosystems.

## Acknowledgments

This work was supported by the National Natural Science Foundation of China (No. 32470388; U22A20558; 32301317), Shandong Provincial Natural Science Foundation (ZR2024QC197), and Tang Scholar Program. The authors thank Professor Jianquan Liu from Lanzhou University and Dr. Kristin Saltonstall from the Smithsonian Tropical Research Institute for providing the geographic information of the known haplotypes.

## Open Research Statement

The raw sequence read data were submitted to the NCBI Sequence Read Archive with the BioProject ID of PRJNA1126092. Data and code are available on Zenodo at https://doi.org/10.5281/zenodo.20362090.

## Author Contributions

L.L. (Lele Liu) led the conceptualization, investigation, methodology, validation, visualization, and writing of the original draft. L.L. also contributed equally to formal analysis and acquired funding. W.S. contributed equally to formal analysis and writing – review & editing, with supporting efforts in investigation. Y.W. contributed equally to conceptualization and funding acquisition, and led the writing – review & editing, with supporting roles in formal analysis and investigation. L.L. (Lele Lin) contributed to conceptualization, funding acquisition, investigation, and writing – review & editing. C.W. contributed to conceptualization, methodology, visualization, and writing – review & editing. H.S. contributed to conceptualization, funding acquisition, investigation, and writing – review & editing. Y.G. contributed equally to writing – review & editing, with supporting roles in investigation, methodology, and visualization. W.G. contributed equally to conceptualization, funding acquisition, and writing – review & editing, and provided overall supervision for the project. All authors reviewed and approved the final manuscript.

