## Supplementary material for "Asymmetric introgression and thermal advantage jointly drive climate-mediated lineage turnover in a mixed-ploidy reed": S

Supporting Information

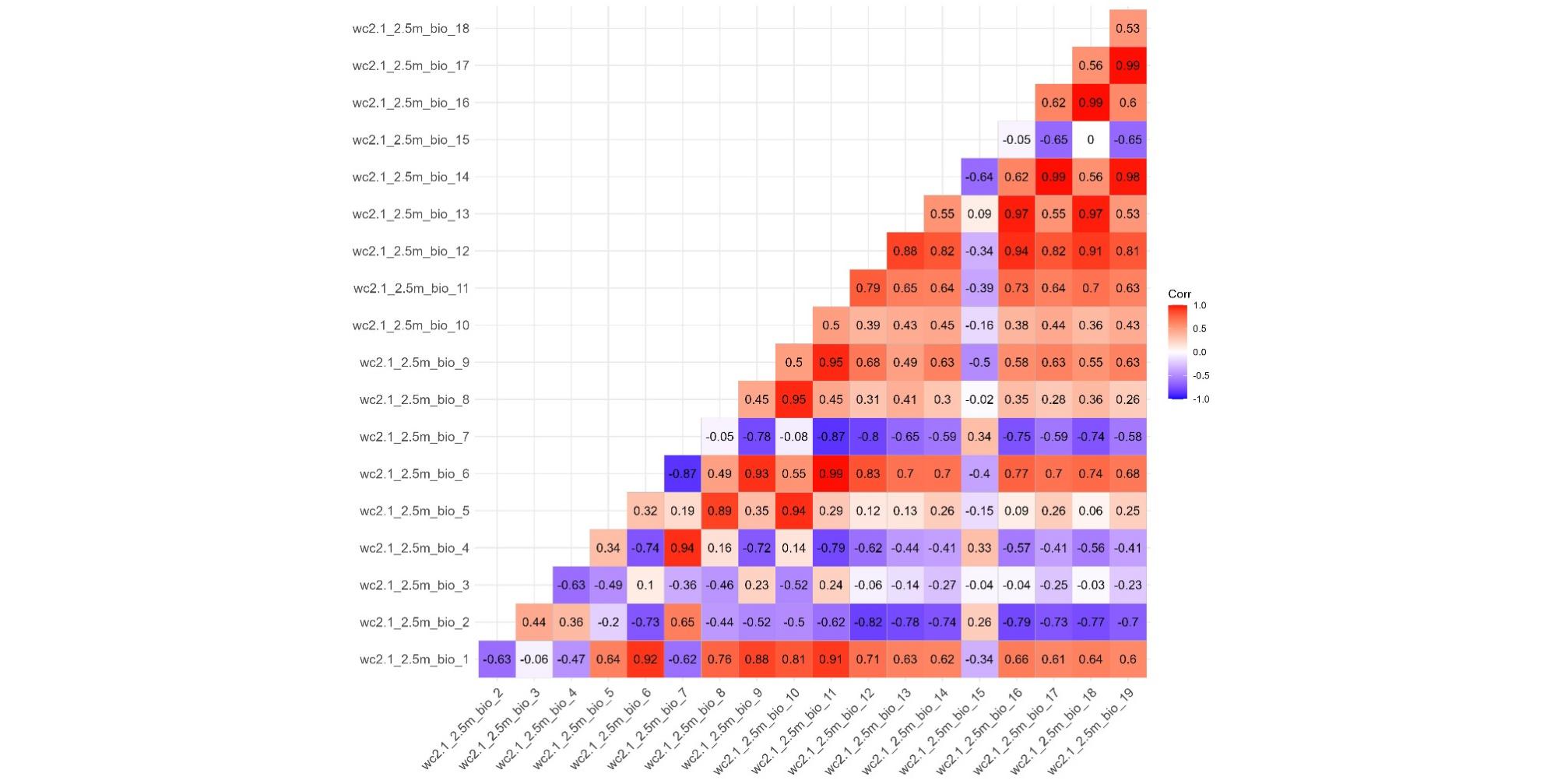

**Figure S1** A correlation heatmap of 19 bioclimatic variables derived from global climate data at 2.5 arc-minute resolution. Values represent Pearson correlation coefficients between variable pairs. Climate data were extracted based on spatially thinned species occurrence records to reduce sampling bias. Correlations were calculated from worldwide bioclimatic layers available from the WorldClim database.

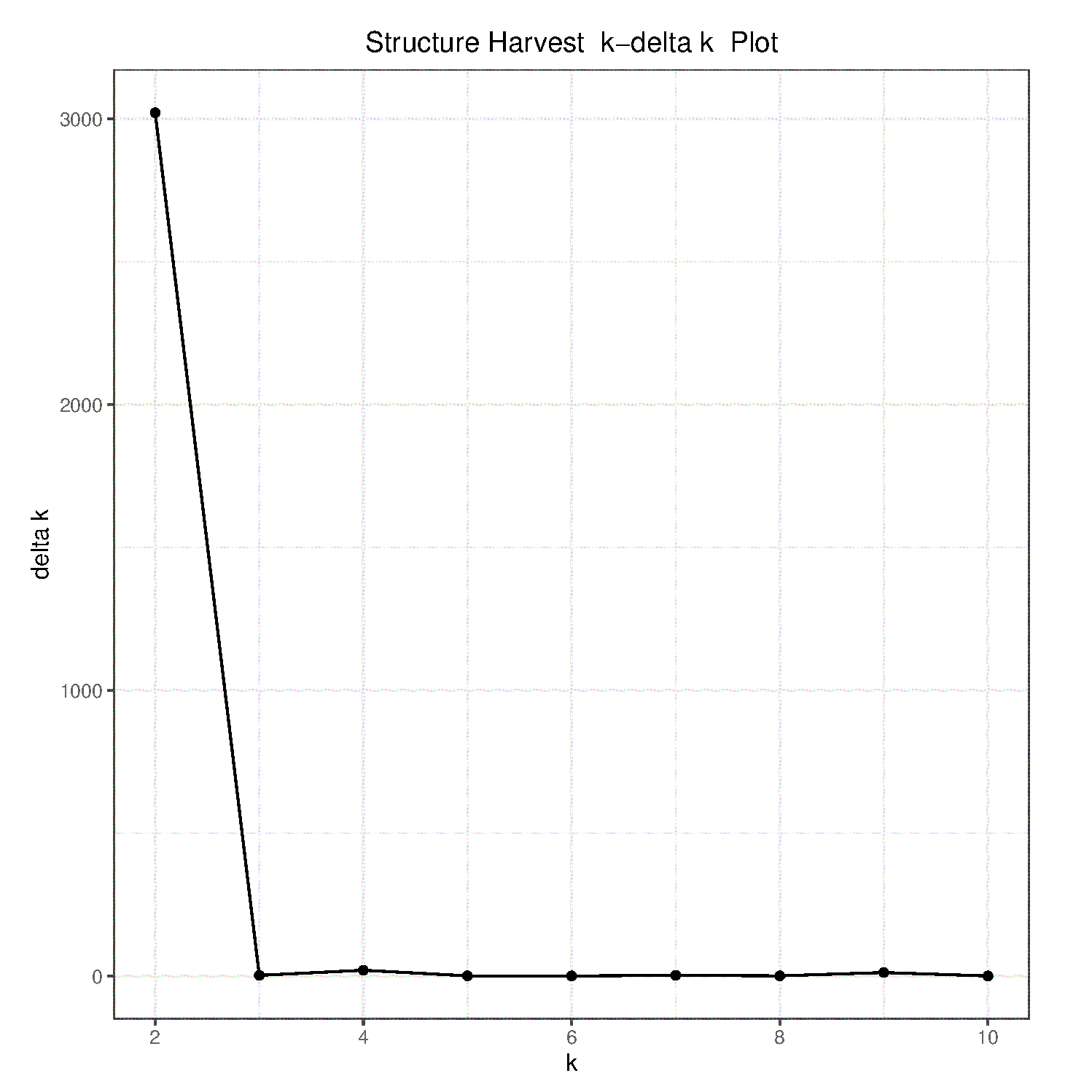

**Figure S2** *k* - delta *k* plot in STRUCTURE analysis. The optimal number of clusters is determined by identifying the *k* value corresponding to the first peak in the delta *k* value.

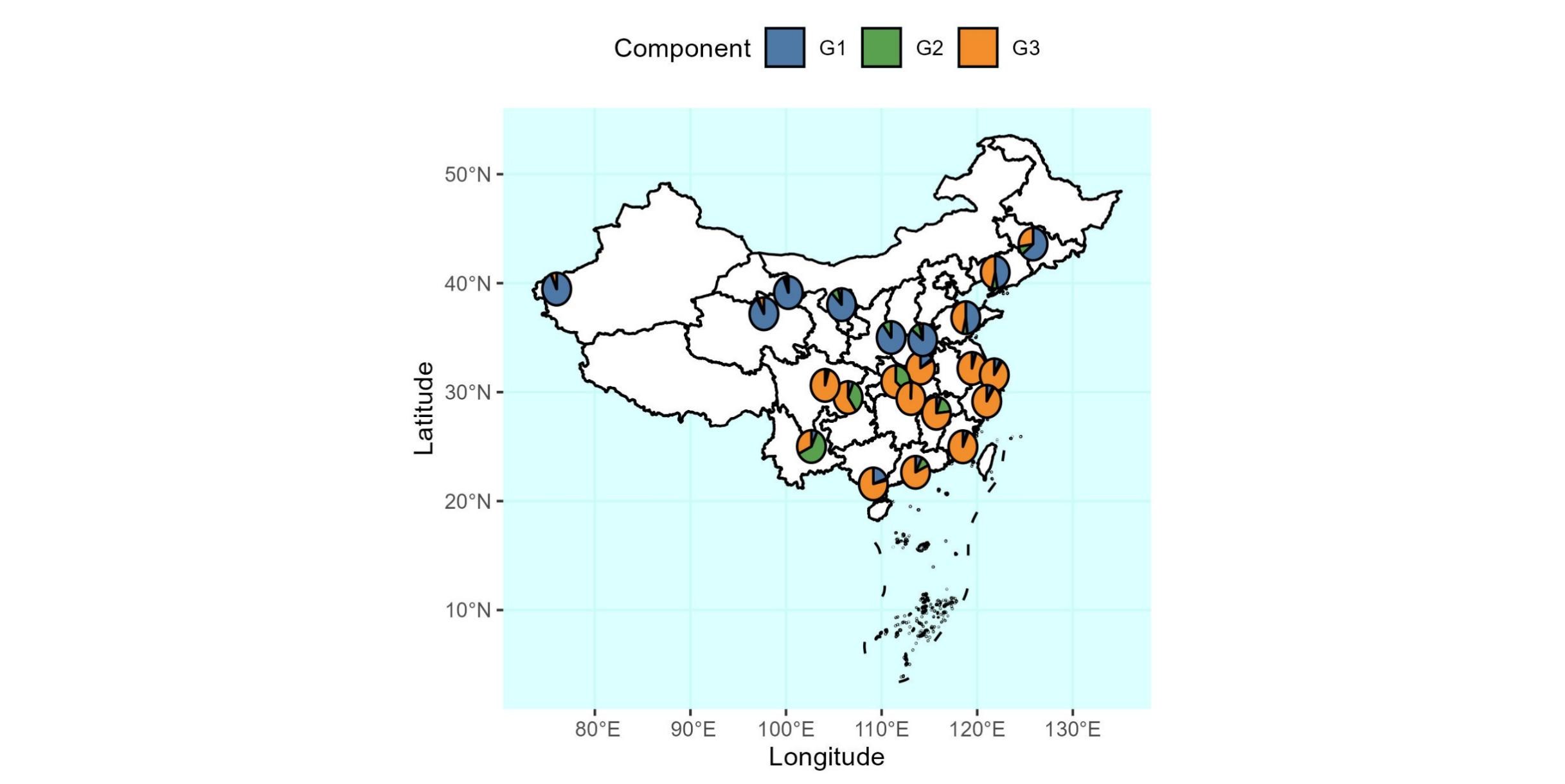

**Figure S3** Population genetic structure of *Phragmites australis* in China (*n* = 458) as inferred from (A) Bayesian clustering analysis of 42 nuclear microsatellites at *k* = 3

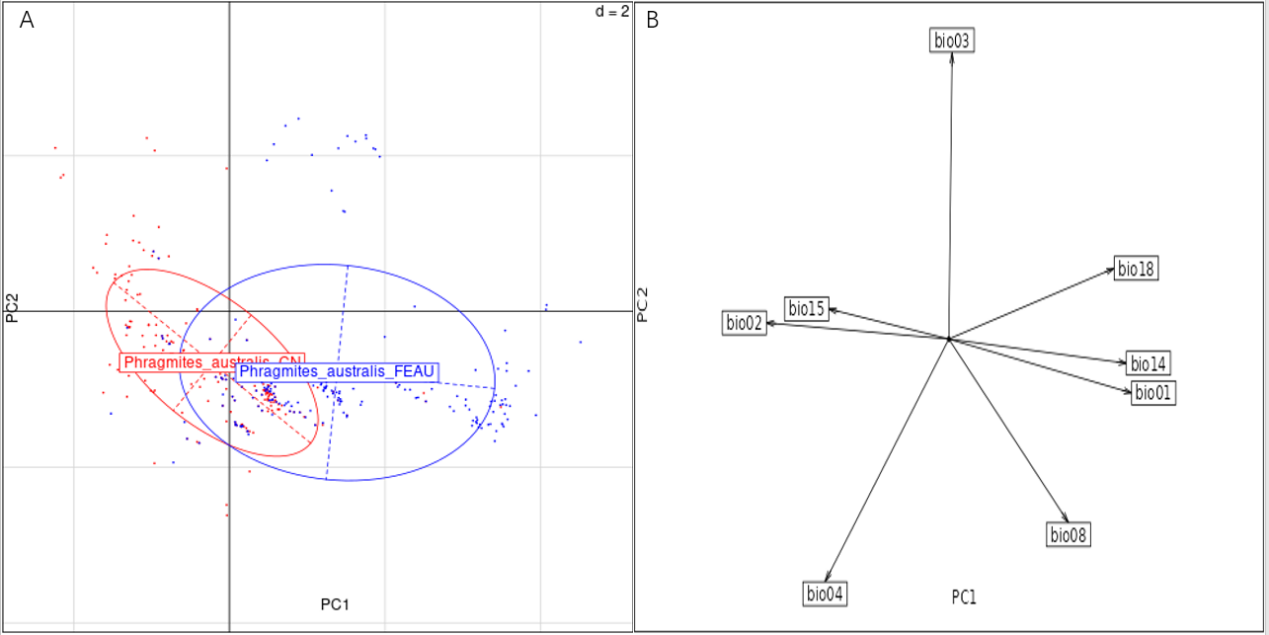

**Figure S4** Principal Components Analysis (PCA) of environmental variables for the CN and FEAU lineages. (A) Scatter plot of occurrences in the PCA space defined by the first two principal components (PC1 and PC2). Points represent individual occurrences, colored by lineage. (B) Loading plot showing the contribution and direction of original bioclimatic variables to PC1 and PC2. Arrows indicate variable loadings, with longer arrows representing stronger contributions to the components.

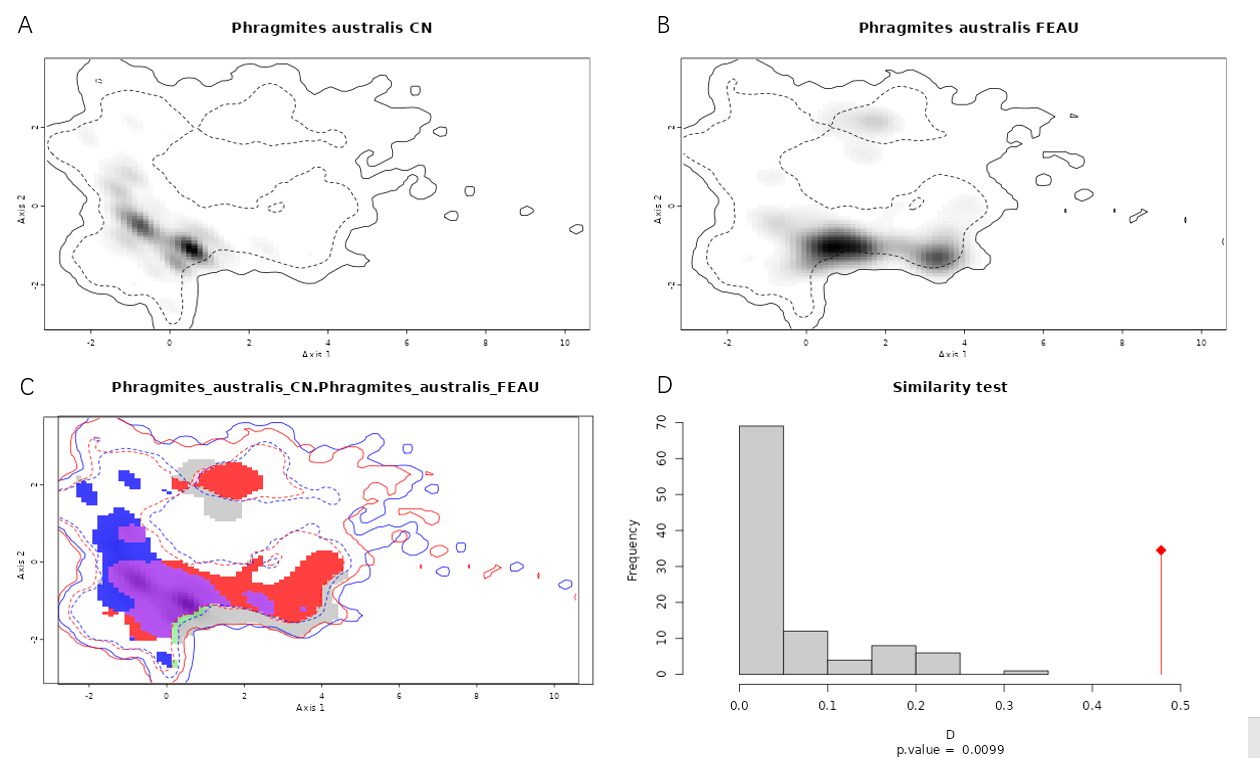

**Figure S5** Niche characterization and overlap between CN and FEAU lineages. (A) and (B) show the gridded density of occurrences in the environmental space, derived from the first two PCA axes, for each lineage. Grey colors indicate higher density of occurrences. (C) Niche overlap in the gridded environmental space, with areas of shared niche space highlighted. (D) Results of niche similarity tests, presenting statistical metrics (D statistics) and significance values indicating the degree of niche conservatism or divergence.

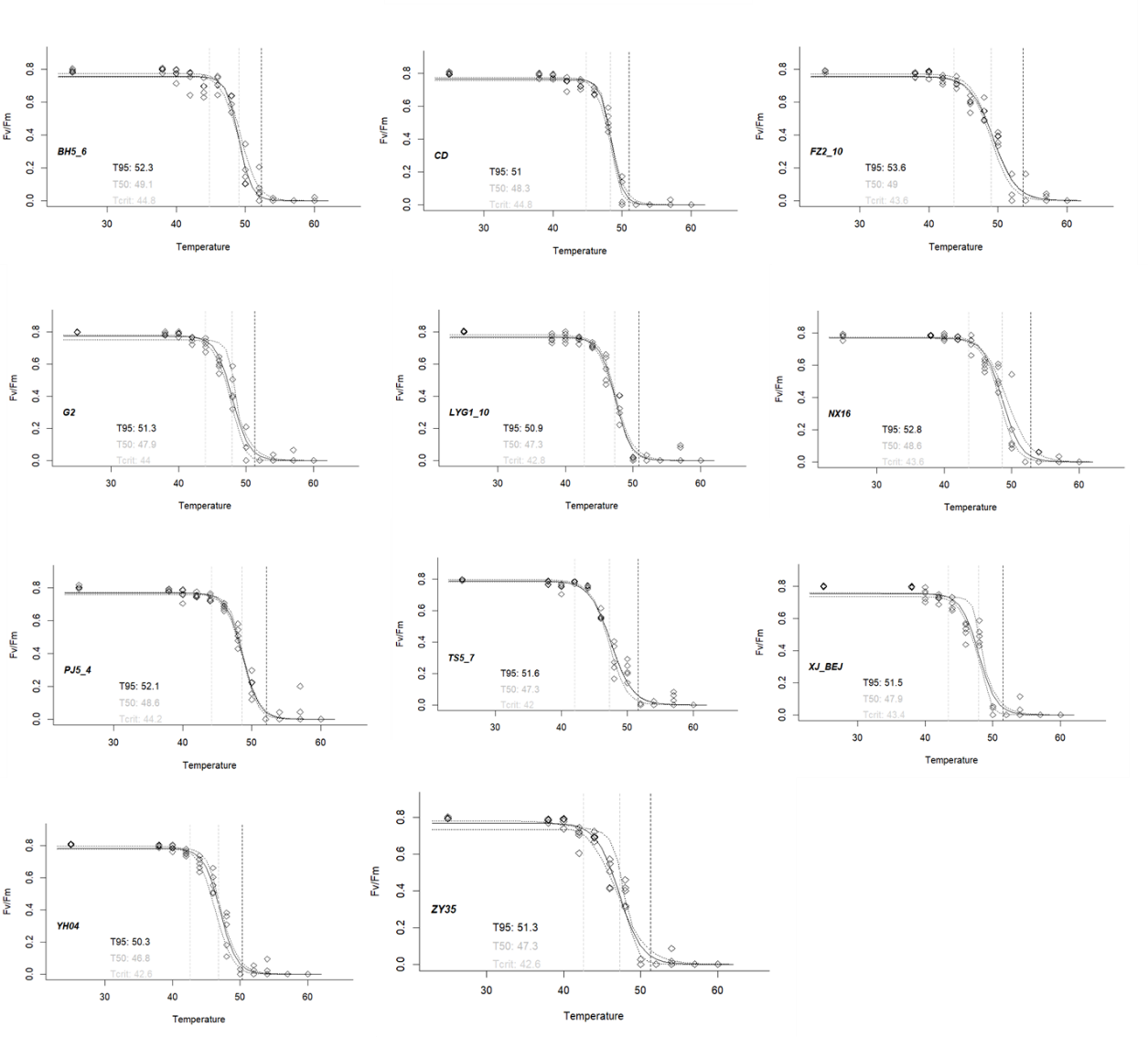

**Figure S6** Representative fitted curve of photosystem II (PSII) heat tolerance for 11 samples, showing the decline in Fv/Fm with increasing temperature. The points represent observed data, the solid line indicates the predicted logistic curve, and the dashed lines show the 95% confidence interval of the fit based on bootstrap resampling. The vertical dashed lines mark the estimated mean critical temperature (*T_crit_*, light gray), the temperature of 50% Fv/Fm reduction (*T_50_,* gray), and the temperature of 95% inhibition (*T_95_*, black), each derived from the model.

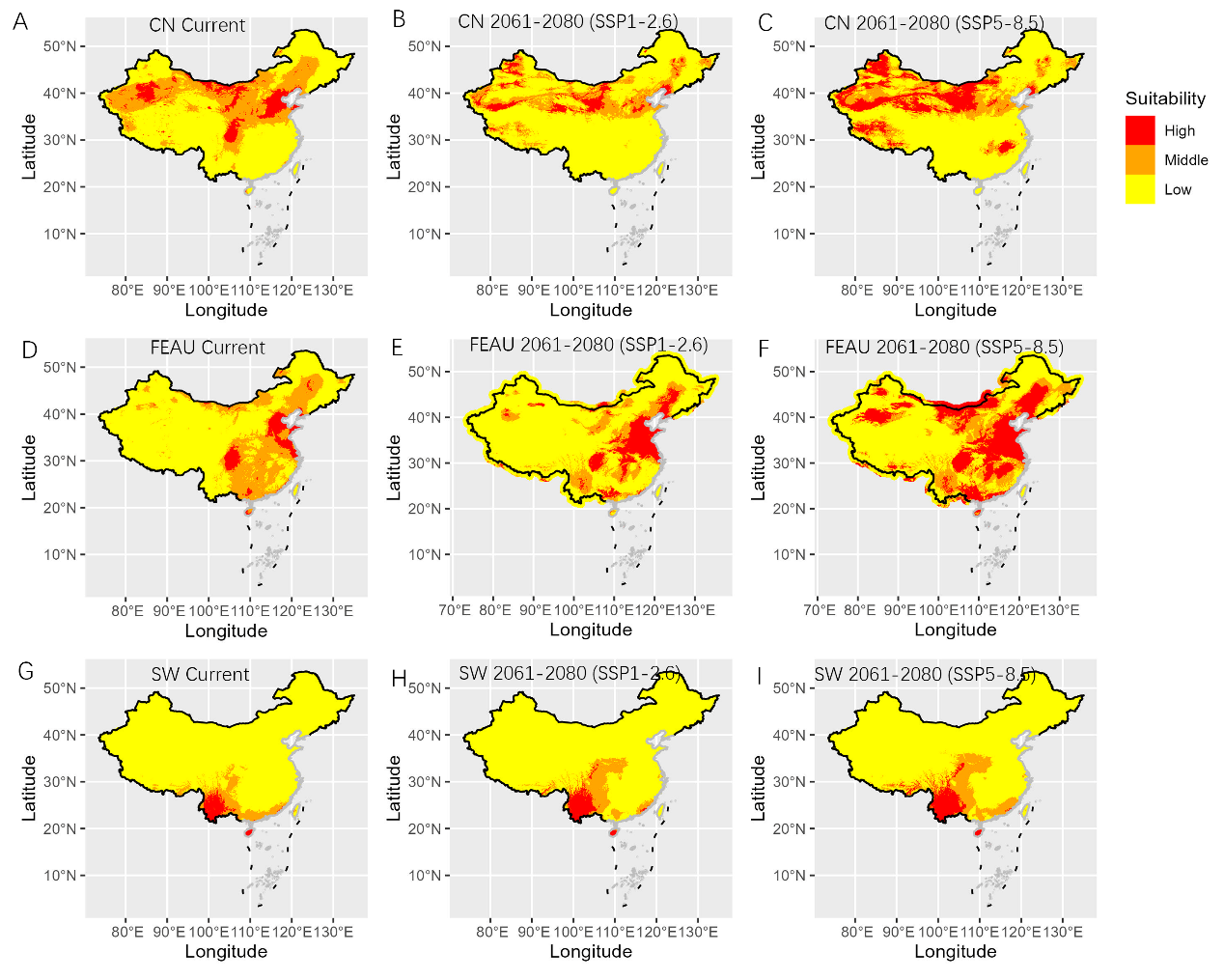

**Figure S7** Predicted potential distribution of three *Phragmites australis* lineages (CN, FEAU, and SW) in China under current and future climate scenarios. The first row (A, B, C) shows the results for the CN lineage, the second row (D, E, F) for the FEAU lineage, and the third row (G, H, I) for the SW lineage. Columns represent different time periods: current distribution (A, D, G), and projected distributions for 2061–2080 under the low-emission scenario SSP1-2.6 (B, E, H) and the high-emission scenario SSP5-8.5 (C, F, I). Predictions were generated using the Maximum Entropy (MaxEnt) model. Suitable habitats are classified into three levels: red indicates high suitability, followed by orange (moderate), and yellow (low).

**Table S1** Collection details of *Phragmites australis* samples used for microsatellite analysis

| **Province** | **Count** | **Longitude (°E)** | **Latitude (°N)** |
| --- | --- | --- | --- |
| Jilin | 14 | 125.8400 | 43.5880 |
| Liaoning | 8 | 121.876 | 41.000 |
| Xinjiang | 11 | 76.0060 | 39.4620 |
| Gansu | 48 | 100.2460 | 39.1370 |
| Ningxia | 26 | 105.8120 | 38.0220 |
| Qinghai | 97 | 97.6860 | 37.1930 |
| Shandong | 68 | 118.8040 | 36.8310 |
| Shanxi | 1 | 110.9970 | 35.0110 |
| Henan | 1 | 114.2960 | 34.8010 |
| Hebei | 39 | 114.0640 | 32.2040 |
| Jiangsu | 23 | 119.4520 | 32.1750 |
| Shanghai | 11 | 121.7850 | 31.5810 |
| Hubei | 5 | 111.4710 | 30.9560 |
| Sichuan | 9 | 104.0870 | 30.6200 |
| Chongqing | 15 | 106.4920 | 29.5000 |
| Hunan | 21 | 113.0650 | 29.4070 |
| Zhejiang | 17 | 120.9990 | 29.1550 |
| Jiangxi | 49 | 115.7290 | 28.0650 |
| Yunan | 10 | 102.6600 | 25.0130 |
| Fujian | 10 | 118.5040 | 24.9840 |
| Guangdong | 7 | 113.5500 | 22.6440 |
| Guangxi | 5 | 109.1340 | 21.5760 |

**Table S2** Characteristics of the microsatellite markers used in this study, including marker name, chromosomal location, quality metrics, sequence, and primer information.

| **Marks** | **Chromosome** | **state** | **motif** | **motif_length** | **target_seq_length** | **primer_F** | **primer_R** |
| --- | --- | --- | --- | --- | --- | --- | --- |
| SSR_3_ | PaChr3 | Good | CA | 2 | 197 | ATTGAATCCACACGTTTCCG | CCATGTGTTAATGTTGTCC |
| SSR_4_ | PaChr18 | Good | CA | 2 | 146 | CAACTCCGTGAATGACATGC | CAGTTTGTGCACTAATGGAC |
| SSR_5_ | PaChr10 | More than 4 alleles (>10 samples) | CA | 2 | 166 | CTTCCTAGGTCAGTATCATCC | GTGGCAGCTGATTGATTTGG |
| SSR_6_ | PaChr12 | Good | TG | 2 | 208 | CTCATGCATCACTTCACAGG | ACACGGACCTAACATCAACC |
| SSR_7_ | PaChr5 | More than 4 alleles (>10 samples) | CA | 2 | 181 | CAAGCATTCTAGTAGTAGC | GTTGCAGCAAGTATTTGG |
| SSR_AU_1128_ | PaChr12 | Good | CGG | 3 | 119 | AGACTCGCCTGCCAGATTTC | CACACTTCTTCTCCCGGCTT |
| SSR_AU_1284_ | PaChr1 | Good | CGC | 3 | 130 | TGCAACCGACGTATGCTTCT | GCCGCGCTTGAAGATAAGGT |
| SSR_AU_151_ | PaChr11 | Good | AGC | 3 | 152 | CGAAGCGTCCCACGAGAG | TTGCTTCTGACGCTACTGGG |
| SSR_AU_1531_ | PaChr22, PaChr10 | More than 4 alleles (>10 samples) | ACA | 3 | 113 | GGAGTGCAAGATGACTGTGC | CCCTGGATCTTGGCCTTCTT |
| SSR_AU_1825_ | PaChr13 | Good | CTG | 3 | 118 | TCAGCCTACTGTTGAAGATGGT | TTCAGTTTCGACTGCCTGCT |
| SSR_AU_1891_ | PaChr4 | Good | AGG | 3 | 103 | GCGTCGATAAGGATGGGGAG | GTCACCCTCTTCAGCCACTC |
| SSR_AU_2008_ | PaChr3 | Good | GCT | 3 | 122 | CGAAACCAAGCGAGCATCAG | AGATCTGCGTCTTCGTCCAC |
| SSR_AU_2013_ | PaChr3 | More than 4 alleles (>10 samples) | GGT | 3 | 108 | GATGGCGGTAGAGGTGACAG | GCCAGAGTATCGGTCACCAC |
| SSR_AU_2160_ | PaChr14 | Good | GTG | 3 | 143 | TGCTTCTGCTCCTCCTGTTC | CCAGTACCAGTACCACCACG |
| SSR_AU_2296_ | PaChr6 | Good | TTC | 3 | 159 | GGAACAGGGCTAATTTCCAAGC | ATAGCCGCAGCCATTAGGAC |
| SSR_AU_2812_ | PaChr13 | Good | AGG | 3 | 115 | CCAGCAAGTGAAGGATCCGA | CTCTCCCCCTTGCACCAAAT |
| SSR_AU_293_ | PaChr16 | Good | CCT | 3 | 157 | AAGCAGCACCCAGATCACTC | TGCTCGCTCCAACAAGACAA |
| SSR_AU_3292_ | PaChr12 | Good | GAG | 3 | 144 | GACCTCTCCGCGTTCTACG | CTGGGGGATCTGCAGGAAG |
| SSR_AU_4223_ | PaChr16 | Good | TTC | 3 | 145 | CGGCCGTAATCCAATCAATGT | AGGAGGAGAAAAAGGGTGCG |
| SSR_AU_4297_ | PaChr3 | Good | AGG | 3 | 198 | ATCGCTTCCTATTGGCCTCG | CTGCACGTCCTCCATCTCC |
| SSR_AU_448_ | PaChr25 | Good | GAA | 3 | 120 | GCCCAAGGTCAGGTTTGGTT | GCCCTTCTCTTCCTCCCTCT |
| SSR_AU_4737_ | PaChr11 | Good | CTC | 3 | 121 | TACGAGCCGACTCCCAGTAA | CTCCTCCTCTCTCTCGGCTT |
| SSR_AU_643_ | PaChr3 | Good | GTC | 3 | 126 | GCCCTCTGCTTAGCTACCAT | TCTCACGCAGGAGGGGAG |
| SSR_AU_656_ | PaChr5 | Failed amplifications (>10 samples) | CAG | 3 | 172 | AGATCACCTCCGATCCTCCC | GTGTCCCCGAATGGTGAGG |
| SSR_CO_15_ | PaChr12 | More than 4 alleles (>10 samples) | CCT | 3 | 113 | CTCCACAAGGTCCTCCACAC | GCCCGTACGGTGATCCAATA |
| SSR_CO_18_ | PaChr15 | Good | TCC | 3 | 126 | GTAGCCTCCCTTGCGGTAG | GAGGAGGAACCGAGGAGGAT |
| SSR_CO_19_ | PaChr19 | More than 4 alleles (>10 samples) | AG | 2 | 116 | ATGTGCGCTTCGAGAATGGA | GGGTTCACGGTGCGTCTG |
| SSR_CO_23_ | PaChr3 | Good | TCT | 3 | 118 | CCTCGACAGGAACCTCCCTA | TAAGAAACCTGCAGGGAGGC |
| SSR_CO_25_ | PaChr8, PaChr14(2), PaChr7(3) | More than 4 alleles (>10 samples) | AG | 2 | 124 | CGGAGGGGTTTATTCCAGGC | AGGGGTGGAGAGGAAGTATCA |
| SSR_CO_27_ | PaChr18 | More than 4 alleles (>10 samples) | TC | 2 | 144 | TTCCCATGTGCTTGTCTCGC | AGATTGAGCAGCCTGTGACG |
| SSR_CO_32_ | PaChr12 | Good | CTAC | 4 | 124 | TGTGTATGCCTGTCACCCAA | ACAAAATCAGCACATCAGCACA |
| SSR_CO_33_ | PaChr6 | Good | TC | 2 | 152 | TCTTCCTCCTCTGGTTCGGT | TGGTACGGTGTTTCGTGGAG |
| SSR_CO_38_ | PaChr6 | Good | GA | 2 | 160 | CAAGCCAAACAAAGCCACCA | TGTGAGGCTCCATGTGCTTT |
| SSR_CO_42_ | PaChr11 | Good | AAGA | 4 | 128 | GGACCTCTTGTCTGAAACCCA | TCGCAAAACAGAAAAGGAGGC |
| SSR_CO_50_ | PaChr23 | Good | AGG | 3 | 108 | GTGCTGGGGGAAGGAGTTC | CTCTCCTCCCTCCACCTCC |
| SSR_CO_69_ | PaChr12 | Good | CTC | 3 | 105 | CCTCCTCCTCTCCCATCTCC | AGAAGGCTAAGGGCAAGGAC |
| SSR_CO_7_ | PaChr20 | Good | TG | 2 | 124 | TGTAGTAATTTGTCGGTTTGCCA | CAACCGAGCAGTAGCCTTGA |
| SSR_CO_8_ | PaChr4 | Failed amplifications (>10 samples) | TGC | 3 | 138 | CAAGTGGTGGTGGTGCCTC | CATTCCGCGATTCTGCCAC |
| SSR_CO_9_ | PaChr3 | Good | CCA | 3 | 163 | CGAGGCCTCCGACTACATC | ACAACGAATGCACGAATCGC |
| SSR_EU_1202_ | PaChr20 | Good | GCG | 3 | 101 | CTTCTCTGCATCGCTTTGCC | ATCTCCACTGCCCTCATCCT |
| SSR_EU_1624_ | PaChr1 | Good | CTC | 3 | 179 | GCTTCGTCTCGCCTAAACCT | CTTCTCGAAGGTGGACTCCG |
| SSR_EU_1667_ | PaChr7 | Good | TCC | 3 | 158 | GCGGTCTAGCTGTTGGATGT | CACAGCGACCAAGCAAAGAG |
| SSR_EU_2523_ | PaChr6 | More than 4 alleles (>10 samples) | GTG | 3 | 129 | ACAGTGGCCGAGAACATCAG | CTGCAACACAACACGGTGAG |
| SSR_EU_2685_ | PaChr14 | More than 4 alleles (>10 samples) | TTC | 3 | 152 | CTCAGCCGCGTCTTCTTCTT | CAACAACACCACACCGACAA |
| SSR_EU_2709_ | PaChr12, PaChr14 | Good | GCA | 3 | 168 | GTGCAGGTGGCCGATGAT | AATGCGACCACACCTCTAGC |
| SSR_EU_3127_ | PaChr9, PaChr12 | More than 4 alleles (>10 samples) | GAG | 3 | 146 | ATGGCAATGGAACGGAGAGG | AACATCCCAAGCGTAGGTGG |
| SSR_EU_3391_ | PaChr23 | Good | GCG | 3 | 128 | CCGCCACCAACTCTTTCTCT | GTAGCTCTTGGGGTGCGAG |
| SSR_EU_3553_ | PaChr2 | Failed amplifications (>10 samples) | ACC | 3 | 150 | GAGTGCAACCCCAACTCCA | GGAGATGTTGCCGAAGACCT |
| SSR_EU_3761_ | PaChr19, PaChr1, PaChr2(2), PaChr6, PaChr18 | Failed amplifications (>10 samples) | CTC | 3 | 175 | AGGAAGCGGAACCCCCTG | CGAAGGACAAGGAGGAGCAG |
| SSR_EU_3780_ | PaChr15 | Good | CTG | 3 | 113 | TAGCAGCTGCAAACCCATGA | ACGAGTTGTTGCTGAGGAGG |
| SSR_EU_4001_ | PaChr4 | Good | GCT | 3 | 109 | AGCTTTCAGACTGTCATGGTCA | GCAGCACAGAAAGGTCATGC |
| SSR_EU_4063_ | PaChr18 | Good | CGT | 3 | 176 | TAGGGGCAGTCGTCGATCTT | AGCTTTACCTTCGTGGCTCC |
| SSR_EU_4112_ | PaChr5 | Good | GAC | 3 | 216 | ACCGTGAAACACCCCATCTC | GGAAAGGTCGTACTCGTCGG |
| SSR_EU_4515_ | PaChr3 | Good | GGA | 3 | 101 | ATCTGTCGGTCGTTCGTTCG | GTATTGCGTGGCCTAGGGTT |
| SSR_EU_643_ | PaChr8 | Good | ATG | 3 | 190 | TGTGGCATTCCAGTAGCAGT | CCATCATCGTCTTCTTTGGCAC |
| SSR_EU_811_ | PaChr5 | Good | GCG | 3 | 179 | AGGAGGGATCCGGAGTTCAA | GGAGTTGAGCTTGACGACGA |
| SSR_EU_914_ | PaChr6 | Failed amplifications (>10 samples) | GCT | 3 | 142 | CCACCTCACCTCTTGGCAAT | CACGAGAGGGTGGTGTGTTT |
| SSR_EU_997_ | PaChr2 | Good | TTG | 3 | 114 | CATCCCTAGTCTATCCCAACTTGT | GCAACAGGACCGGTCACATA |

**Table S3** *Phragmites australis* samples used in the common garden experiment, detailing their designated name, haplotype, and geographical origin.

| **ID** | **Garden** | **Group** | **Haplotype** | **Latitude** (°N) | **Longitude** (°E) |
| --- | --- | --- | --- | --- | --- |
| YRD9 | Jinan-Panjin | CN | O | 38.0037 | 118.9496 |
| YRD8 | Jinan-Panjin | CN | O | 37.9719 | 118.8778 |
| YRD7 | Jinan-Panjin | CN | O | 37.8447 | 118.9930 |
| YRD5 | Jinan-Panjin | CN | O | 37.8338 | 119.0382 |
| YRD4 | Jinan-Panjin | CN | O | 37.7950 | 118.8973 |
| YRD3 | Jinan-Panjin | CN | O | 37.7458 | 119.1342 |
| YRD2 | Jinan-Panjin | CN | O | 37.7458 | 119.1342 |
| Salt1 | Jinan-Panjin | CN | O | 38.0458 | 118.6777 |
| DW1 | Jinan-Panjin | FEAU | P | 37.7929 | 118.4386 |
| CN2028 | Jinan-Panjin | CN | O | 41.2069 | 122.0279 |
| CN2026 | Jinan-Panjin | FEAU | P | 41.2069 | 122.0279 |
| CN2024 | Jinan-Panjin | FEAU | P | 41.0892 | 122.0578 |
| CN2023 | Jinan-Panjin | FEAU | P | 36.4551 | 120.6805 |
| CN2021 | Jinan-Panjin | FEAU | P | 37.0676 | 118.0605 |
| CN2020 | Jinan-Panjin | FEAU | P | 37.0712 | 118.0608 |
| CN2019 | Jinan-Panjin | FEAU | P | 37.0712 | 118.0607 |
| CN2018 | Jinan-Panjin | CN | O | 37.0484 | 118.0653 |
| CN2017 | Jinan-Panjin | CN | O | 37.0484 | 118.0653 |
| CN2016 | Jinan-Panjin | FEAU | P | 30.2593 | 120.0623 |
| CN2014 | Jinan-Panjin | FEAU | P | 30.1652 | 120.0942 |
| DY3_10 | Qingdao-Shanghai | FEAU | P | 37.8306 | 119.0783 |
| LANZHOU | Qingdao-Shanghai | CN | O | 36.2049 | 103.7024 |
| PJ1_4 | Qingdao-Shanghai | FEAU | P | 40.8977 | 121.7893 |
| QH05 | Qingdao-Shanghai | CN | M | 35.8456 | 102.8543 |
| SH1_8 | Qingdao-Shanghai | FEAU | P | 31.6844 | 121.6522 |
| TS1_3 | Qingdao-Shanghai | CN | O | 39.0254 | 118.3356 |
| TS2_1 | Qingdao-Shanghai | CN | O | 39.0414 | 118.2834 |
| TS3_1 | Qingdao-Shanghai | CN | O | 39.0613 | 118.2254 |
| TS4_3 | Qingdao-Shanghai | CN | O | 39.0774 | 118.2062 |
| TS5_7 | Qingdao-Shanghai | CN | O | 39.0987 | 118.1895 |
| WZ1_5 | Qingdao-Shanghai | FEAU | P | 28.0054 | 120.7837 |
| WZ4_7 | Qingdao-Shanghai | FEAU | P | 27.9836 | 120.7745 |
| WZ5_10 | Qingdao-Shanghai | FEAU | P | 27.9852 | 120.7671 |
| XJ_BEJ | Qingdao-Shanghai | CN | M | 39.4616 | 76.0062 |
| YC4_4 | Qingdao-Shanghai | FEAU | P | 33.6182 | 120.4933 |
| YC5_10 | Qingdao-Shanghai | FEAU | P | 33.6242 | 120.4817 |
| YUNCHENG | Qingdao-Shanghai | CN | O | 35.0106 | 110.9972 |

**Table S4** Heat tolerance metrics of different *Phragmites australis* genotypes, derived from chlorophyll fluorescence curve analysis.

|  | **Group** | ***T_crit_*** | ***T_50_*** | ***T_95_*** |
| --- | --- | --- | --- | --- |
| G2 | CN | 43.8 | 47.9 | 51.3 |
| NX16 | CN | 43.4 | 48.5 | 52.7 |
| TS5_7 | CN | 41.8 | 47.6 | 52.5 |
| XJ_BEJ | CN | 44.2 | 47.9 | 50.8 |
| YH04 | CN | 42.4 | 47 | 50.8 |
| ZY35 | CN | 41.8 | 47.1 | 51.5 |
| BH5_6 | FEAU | 45.6 | 49 | 51.8 |
| CD | FEAU | 44.4 | 48.3 | 51.2 |
| FZ2_10 | FEAU | 43.4 | 49.3 | 54.3 |
| LYG1_10 | FEAU | 43.2 | 47.3 | 50.8 |
| PJ5_4 | FEAU | 44.4 | 48.7 | 52.2 |
| YN01 | FEAU | 40.4 | 46.6 | 51.8 |

**Table S5** Best-performing parameter combinations for the species distribution models of different *Phragmites australis* lineages.

|  | **FEAU** | **CN** | **SW** |
| --- | --- | --- | --- |
| fc | LQH | LQHP | LQ |
| rm | 0.5 | 0.5 | 1 |
| tune.args | fc.LQH_rm.0.5 | fc.LQHP_rm.0.5 | fc.LQ_rm.1 |
| auc.train | 0.945856 | 0.946841 | 0.964724 |
| cbi.train | 0.928 | 0.973 | 0.858 |
| auc.diff.avg | 0.057 | 0.082 | 0.078 |
| auc.diff.sd | 0.061 | 0.037 | 0.079 |
| auc.val.avg | 0.898 | 0.869 | 0.936 |
| auc.val.sd | 0.068 | 0.039 | 0.078 |
| cbi.val.avg | 0.757 | 0.731 | 0.517 |
| cbi.val.sd | 0.173 | 0.154 | 0.187 |
| or.10p.avg | 0.190 | 0.283 | 0.250 |
| or.10p.sd | 0.139 | 0.137 | 0.500 |
| or.mtp.avg | 0.056 | 0.046 | 0.250 |
| or.mtp.sd | 0.112 | 0.054 | 0.500 |
| AICc | 4446.910 | 4477.249 | 603.704 |
| delta.AICc | 0 | 0 | 0 |
| w.AIC | 0.776 | 1.000 | 0.658 |
| ncoef | 47 | 48 | 7 |

**Table S6** Projected changes in suitable habitat distribution (%) for three *Phragmites australis* lineages in China under current and future climate scenarios (2061–2080).

| Lineage | Stage | High (%) | Middle (%) | Low (%) |
| --- | --- | --- | --- | --- |
| CN | Current | 9.4 | 28.5 | 62.1 |
| CN | 2061-2080 (SSP1-2.6) | 6.6 | 18.6 | 74.8 |
| CN | 2061-2080 (SSP5-8.5) | 16.5 | 20.0 | 63.4 |
| FEAU | Current | 6.6 | 24.5 | 69.0 |
| FEAU | 2061-2080 (SSP1-2.6) | 10.9 | 14.5 | 74.6 |
| FEAU | 2061-2080 (SSP5-8.5) | 25.2 | 18.1 | 56.7 |
| SW | Current | 3.3 | 6.6 | 90.1 |
| SW | 2061-2080 (SSP1-2.6) | 4.1 | 8.7 | 87.2 |
| SW | 2061-2080 (SSP5-8.5) | 5.3 | 10.7 | 83.9 |
